# Monocyte Lineage Expansion Drives Transcriptomic Individuality in Genetically Identical Armadillo Quadruplets

**DOI:** 10.64898/2026.03.30.715171

**Authors:** Risa Karakida Kawaguchi, Sara Ballouz, Maria T. Pena, Leon French, Frank M. Knight, Linda B. Adams, Jesse Gillis

**Affiliations:** Cold Spring Harbor Laboratory, Cold Spring Harbor, NY, 11724, USA; Center for iPS Cell Research and Application, Kyoto University, Kyoto, Japan; Graduate School of Pharmaceutical Science, The University of Tokyo, Tokyo, Japan; School of Computer Science and Engineering, Faculty of Engineering, University of New South Wales Sydney, Sydney, NSW, Australia; United States Department of Health and Human Services, Health Resources and Services Administration, Health Systems Bureau, National Hansen’s Disease Program, Baton Rouge, LA, USA; Department of Physiology and Donnelly Centre for Cellular and Biomolecular Research, University of Toronto, Toronto, ON, Canada; University of the Ozarks, Clarksville, AR, 72830, USA; Former United States Department of Health and Human Services, Health Resources and Services Administration, Health Systems Bureau, National Hansen’s Disease Program, Baton Rouge, LA, USA

**Author notes:** These authors contributed equally to this work (Risa Karakida Kawaguchi, Sara Ballouz).

## Abstract

Genetic diversity shapes phenotypes, yet even genetically identical individuals differ. In nine-banded armadillo (*Dasypus novemcinctus*) quadruplets, we previously showed that allele-specific expression (ASE) imbalances provide a stable molecular fingerprint of individuality. Here, we test whether such transcriptomic individuality reflects functional biological differences. We profiled bulk blood RNA from five cohorts of genetically identical quadruplets across three time points, and found persistent gene-expression signatures that predict individual identity. Focusing on a highly variable cohort, we then performed single-cell RNA-seq and ATAC-seq. In this litter, the most stably distinct individual showed an expanded monocyte-lineage compartment and gene-expression programs enriched for inflammatory and differentiation pathways. These cell-type and regulatory differences were stable over time and robust to experimental leprosy infection. Together, our results link transcriptomic individuality to lasting differences in immune-cell composition, illustrating how early stochastic events can produce persistent, biologically meaningful divergence among genetically identical individuals.

**Teaser:** Genetically identical armadillo quadruplets are identifiable by transcriptomic signatures shaped by cellular composition variation.

## Introduction

Genetically identical individuals raised in similar environments can nevertheless diverge markedly in their phenotypes (*1–4*). Understanding the origins of this non-genetic individuality is challenging because it reflects influences that fall outside both genotype and measurable environmental exposures. One plausible contributor is stochasticity in developmental processes, meaning random molecular or cellular events whose consequences can propagate across scales (*5*). A useful analogy comes from Waddington’s depiction of cell-fate decisions as trajectories shaped by an underlying “epigenetic landscape” (*6*). Although originally formulated for cellular differentiation, this idea illustrates how small early perturbations could, in principle, bias organism-level outcomes. Experimental systems support this view, showing that noise in gene expression or protein abundance can generate persistent divergence among genetically identical cells or organisms (*7*, *8*). Random monoallelic expression offers another potential source of such divergence (*9*). In previous work on *Dasypus novemcinctus* (the nine-banded armadillo), we found that stable allele-specific expression (ASE) imbalance patterns distinguish genetically identical siblings, indicating that early stochastic events can leave persistent epigenetic signatures (*10*).

It is not known whether the transcriptomic individuality we previously described corresponds to stable differences in biological function. In general, much molecular variation is expected to be neutral, and many allele-specific expression (ASE) imbalances likely fall into this category, meaning they can persist as individual markers without altering total gene output or phenotype. Total expression of a gene (from both alleles), in contrast, has more direct links to cellular physiology, although it is more dynamic and sensitive to environmental influences. Several routes could, in principle, connect non-genetic individuality to functional differences. One possibility is that early stochastic events bias the relative abundance of specific cell types, producing persistent but subtle differences in cellular composition. Another possibility is that individuals differ primarily in the activation state or regulatory programs of the same cell types, leading to stable differences in transcriptional tone without changes in lineage proportions. Clonal haematopoiesis in aging humans illustrates how small lineage or state biases can have long-term consequences for immune function (*11*). Similar non-heritable processes could contribute to stable individuality among genetically identical siblings.

In this study, we ask whether transcriptomic individuality extends beyond neutral molecular signatures to include stable functional differences. We analyzed bulk blood transcriptomes from five sets of genetically identical armadillo quadruplets raised under controlled conditions, and found that each sibling maintained a consistent gene expression profile across three sampling points. These profiles contain thousands of genes whose ranked expression distinguishes individuals despite identical genotype and shared environment. To test whether these differences reflect underlying physiology, we generated single-cell RNA-seq and single-cell ATAC-seq data from one litter that showed particularly high phenotypic and transcriptomic variability. In this cohort, the most distinctive sibling displayed a clear shift in monocyte-lineage abundance together with an associated innate immune expression program, whereas its siblings showed more modest lineage- or state-specific differences. These findings link transcriptomic individuality to coherent differences in cellular composition and regulatory activity. They also illustrate how early stochastic influences can give rise to lasting, biologically meaningful individuality among genetically identical animals.

## Results

### Bulk Transcriptome Analysis of Genetically Identical Quadruplets Reveals Structured Phenotypic Variability

Under controlled conditions, genetically identical individuals might generally be expected to show limited phenotypic divergence. In the five armadillo quadruplet litters we examined (20 animals in total, each litter derived from a single embryo), we nevertheless observed consistent differences in basic physiological traits (Fig. 1A). We quantified within-litter variability using the coefficient of variation for measures such as body weight and white blood cell counts obtained from clinical chemistry and complete blood count (CBC) assays. These traits differed in a structured way across litters. Some groups showed little dispersion, whereas others were noticeably more variable. The 16- 90 litter, for example, had one of the highest coefficients of variation in body weight and the greatest variability in white blood cell counts (Fig. 1B-E). These observations indicate that factors other than genotype and shared environment contribute to stable phenotypic differences among siblings.

**Fig. 1.**
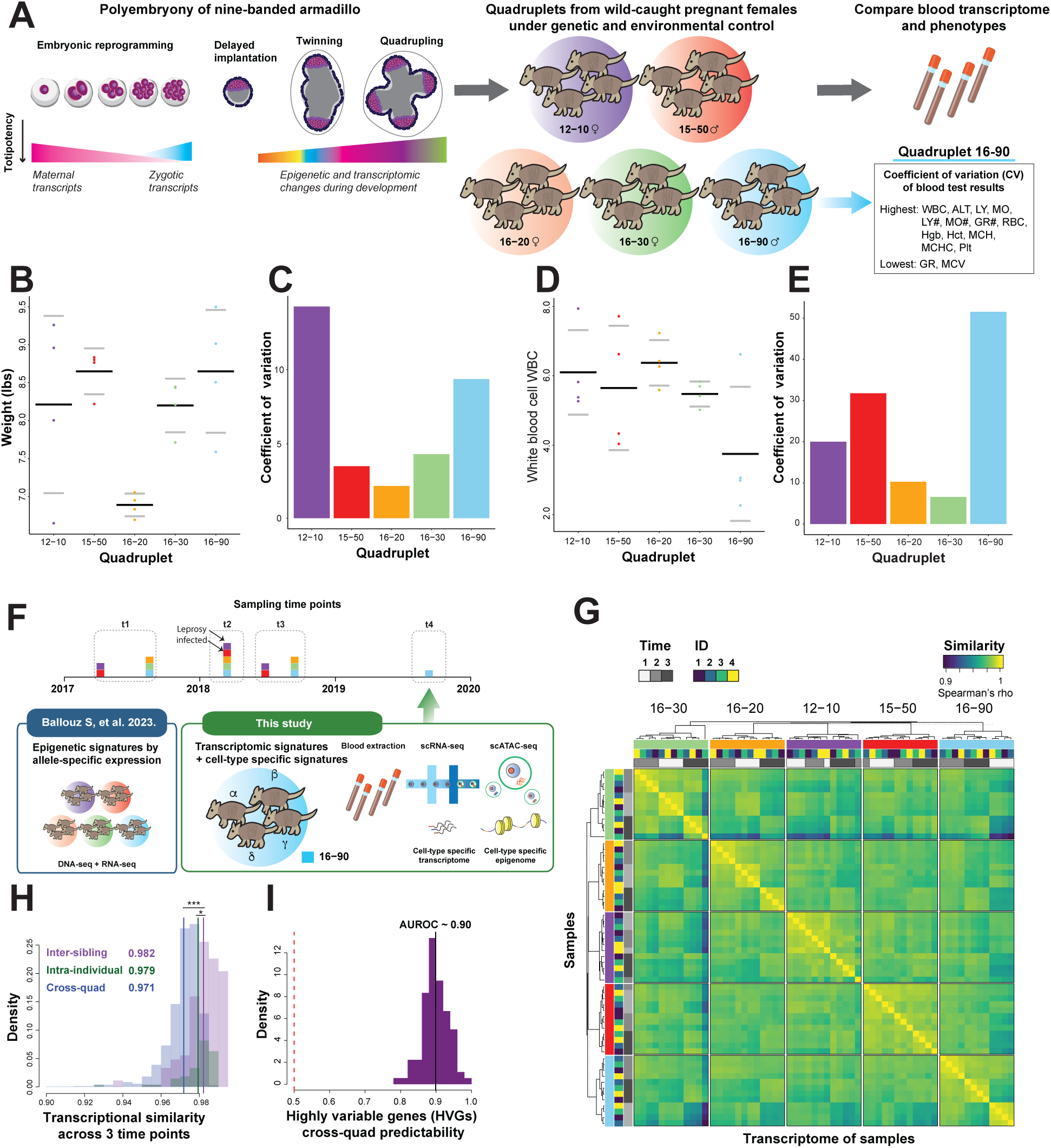
Dissecting non-genetic variation in armadillo sibling cohort data. (**A**) Polyembryony in the nine-banded armadillo enables efficient sampling of genetically identical siblings under environmental conditions. Quadruplets from five different lineages were raised in the same colony at the laboratory, and blood samples were collected for transcriptomic analysis and phenotypic assessment. Quadruplet group 16-90 shows the highest phenotypic variation across 13 blood-based traits. (**B-E**) Phenotypic variations of each quadruplet group: 1) the weight distribution and its coefficient of variation (**B**) and (**C**), and 2) white blood cell (WBC) count (×10³ cells/mm³) for individuals from five quadruplets (**D**) and (**E**). (**F**) Timeline depicting previously collected blood samples and newly obtained samples used for scRNA-seq and scATAC-seq analyses of quadruplet 16-90. (**G**) Correlation matrix of bulk gene expression profiles for five quadruplet sets at three time points. (**H**) Distributions of correlations from (**G**) categorized as intra-individual across time (intra-ind), within quadruplet sets (inter-ind), and across quadruplet sets (cross-quad). (**I**) AUROC distribution from leave-one-out analysis to evaluate the consistency of highly variable genes (HVGs) across quadruplet groups.

We next asked whether transcriptomic profiles showed evidence of the same structured individuality. Bulk RNA sequencing of peripheral blood mononuclear cells was performed for all five quadruplet sets at three time points over roughly one year (*10*) (Fig. 1F). Gene expression correlations confirmed that each sibling maintained a highly stable transcriptome across time and that these within-individual profiles were more similar to one another than to profiles from unrelated animals (Fig. 1G). Intra-individual correlations were uniformly high (median Spearman ρ = 0.979) and close in magnitude to correlations among siblings, although the small difference between these two groups was statistically significant (ρ = 0.982 on average for sibling pairs, p = 0.008; Fig. 1H). To place these values in context, we compared log fold changes of homologous genes in armadillos to data from human monozygotic and dizygotic twins. Armadillo siblings showed substantially lower transcriptomic dispersion than human monozygotic twins (Fig. S1), a result consistent with reduced environmental variation in the controlled armadillo colony relative to human cohorts.

### Highly Variable Genes Within Sibling Cohorts Are Shared Across Cohorts

We next asked whether certain genes contributed more strongly than others to the observed individuality. If transcriptomic differences among siblings were purely diffuse noise, we would not expect the same genes to vary consistently within different litters. For each litter at each time point, we ranked genes by their coefficient of variation across siblings and selected the top 100 highly variable genes. We then evaluated how well the HVG sets from one litter predicted the HVGs identified in another by calculating AUROC scores across all cross-litter comparisons (Fig. 1I). These scores were high, with a mean AUROC of 0.90, indicating that the same genes tend to show elevated within-litter variability across independent groups. This pattern is unlikely under a random model and suggests that individuality is partly encoded in a subset of genes that reliably exhibit greater variance among genetically identical siblings.

### Rank-Ordered Gene Expression Patterns Identify Stable Identity Signatures

We next examined whether individuality was reflected not only in variability at single time points but also in the stability of expression differences across time. To do this, we assessed whether the relative ordering of siblings by gene expression, within a litter, was preserved across the three sampling points. For example, if a sibling had the highest expression of a gene at the first time point, we asked whether it tended to remain highest at the later time points (Fig. 2A). This rank-based approach parallels our earlier analysis of allele-specific imbalance and reduces sensitivity to shifts in absolute expression levels.

**Fig. 2.**
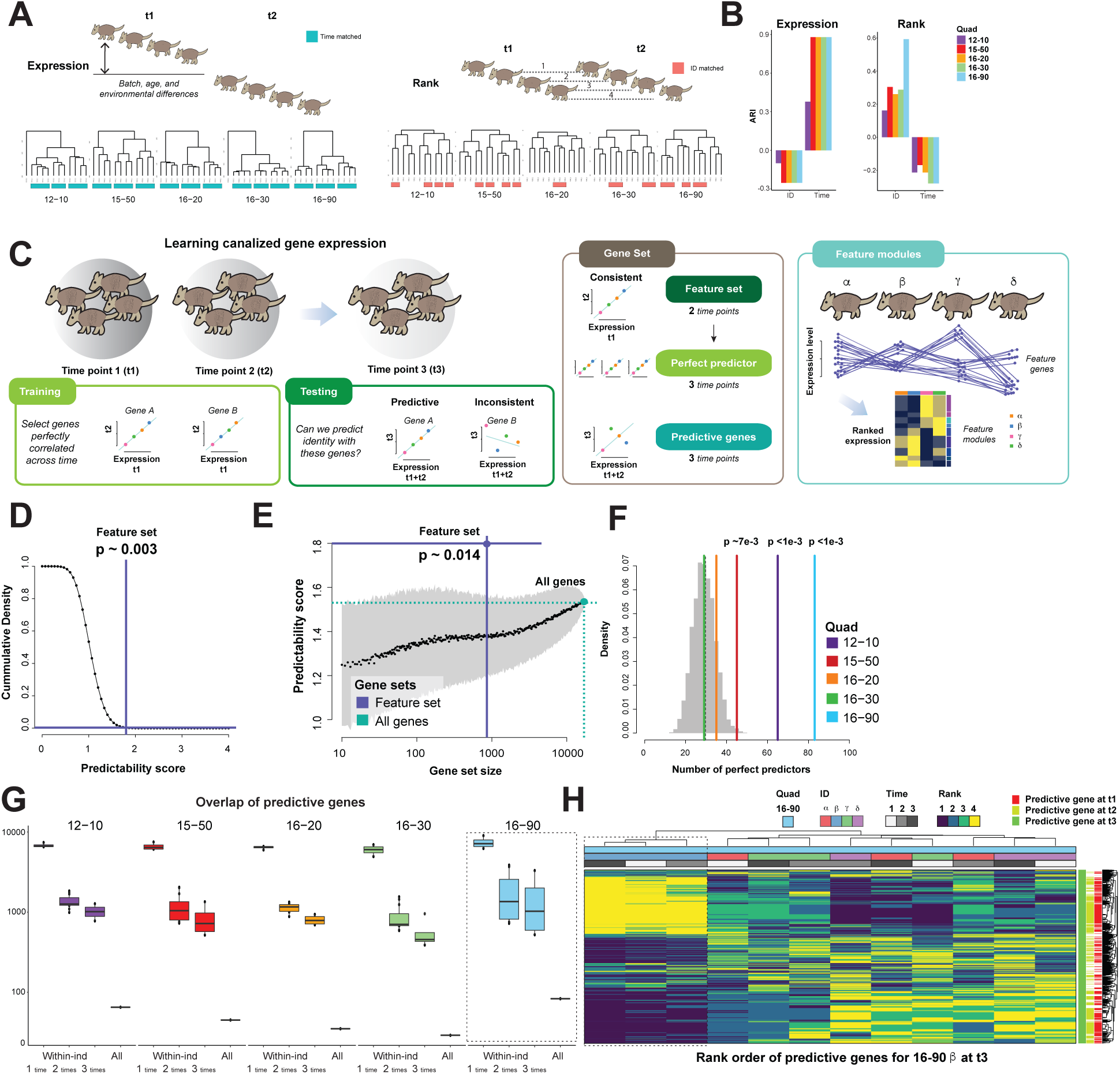
Gene expression-based identity prediction by sibling-specific signatures. (**A**) Clustering of transcriptome profiles among siblings before (left) and after (right) ranking within a given time point (**B**) Adjusted Rand Index (ARI) measuring how well the samples from the same individual (ID) or time point are assigned to the same cluster as shown in (**A**). (**C**) Workflow illustrating the procedure for predicting individual identity and defining identity-predictable genes. A feature set is optimized for the prediction of each individual to make the gene expression ranking consistent between two time points. A perfect predictor can further predict the third point based on the average rank of two time points. For functional analysis, each gene is classified into the feature modules based on the expression patterns. (**D,E**) Analytical (**D**) and empirical (**E**) null of identity predictability scores with the score of feature gene sets 1.8. (**F**) Number of perfect predictors for each quadruplet with p-values computed by random background distribution (gray). (**G**) Number of common predictive genes within the same individual and within the siblings from five lineages. (**H**) Example of ranked gene expression profiles for predictive genes identified at time point t_3_ from the quadruplet group exhibiting the highest predictability (16–90).

Clustering based on raw expression values grouped samples primarily by time point rather than by individual, indicating that global expression levels changed systematically as the armadillos developed and aged over the sampling period (Fig. 2A, left). In contrast, clustering based on rank-ordered expression grouped samples more strongly by individual identity (Fig. 2A, right). Adjusted Rand Index scores confirmed this pattern: rank-based clustering aligned more closely with true individual labels, whereas raw-expression clustering reflected developmental time (Fig. 2B). These results suggest that relative expression ordering captures stable, individual-specific transcriptomic features that are not apparent from absolute expression alone.

To quantify how well gene expression patterns could identify individual siblings, we built a supervised classifier based on ranked expression (Fig. 2C). For each litter, we used two time points as training data and asked whether the ranked expression of selected genes could correctly match each sibling in the remaining time point. Genes were included as features if their ranked expression was consistent between the two training time points. For each gene, we compared the ordering of the four siblings in the training data with their ordering in the held-out time point. A gene contributed one point for each sibling whose position in the ranked order matched between training and test data. Summing across genes yielded a score for each sibling, and the final identity score for a litter was the number of siblings, out of four, that were correctly assigned to themselves on average across all training–test splits.

Across litters, the classifier achieved a mean identity score of 1.8, which is significantly higher than the score of approximately 1.0 expected by chance (Fig. 2D; empirical p-value ∼0.014). Randomly selected gene sets produced much lower scores, and their predictability increased only gradually with gene-set size (Fig. 2E). The strength of the identity signal differed across litters. Two groups, 12-10 and 16-90, had especially strong signatures, each containing more than 60 genes that maintained a perfectly consistent rank order across all three time points (Fig. 2F). These “perfect predictors” did not generally overlap with the highly variable genes identified earlier and were not distinguished by higher average expression (Fig. S2). Stable rank ordering of gene expression therefore identifies individual-specific signatures that differ across litters and are not explained by variability or expression level alone.

### Environmental Robustness of Individuality: The Role of Leprosy Exposure

Two of the five quadruplet groups (12-10 and 15-50) were experimentally infected with *Mycobacterium leprae* after the second time point as part of an unrelated study at the National Hansen’s Disease Program, where the colony is maintained for leprosy research. Although this exposure was not a focus of our experimental design, it provides a natural test of whether individuality signals persist in the presence of physiological disturbance. Even modest immune challenges can reshape blood transcriptomes, so the infection offers a useful opportunity to examine robustness, even if the perturbation is not severe.

To assess the impact of infection, we applied a k-mer based classifier to detect pathogen-derived sequences in the RNA-seq data (*12*). No *M. leprae* sequences were detected, consistent with early or localized infection, and detection of other organisms did not differ by infection status. Differential expression analysis identified some infection-responsive genes, but these did not overlap with established leprosy-associated signatures (Fig. S3). Although the infection was mild, individuality signatures remained clearly detectable in the exposed litters. This suggests that the identity-specific patterns are stable under at least modest immune perturbation, while leaving open the possibility that stronger challenges could produce additional effects.

### Total Expression-Based Individuality is Independent of Allelic Imbalance Signatures

We then asked how the total-expression identity signatures relate to the allele-specific expression (ASE) signatures described in our earlier study (*10*). ASE-based individuality arises from persistent monoallelic biases and can distinguish genetically identical siblings without altering total transcript abundance. Consistent with this expectation, the allelically imbalanced markers identified previously did not show elevated variability in their overall expression across individuals (Fig. S4). These genes therefore maintain stable total expression despite their allelic differences, indicating that the ASE-based and total-expression-based individuality signals represent largely independent axes of variation.

### Broadening Identity Signatures Through Predictive Genes

Perfect predictors are highly stringent, since they require an identical rank order across all three time points, and therefore identify only a limited set of genes. To capture weaker but still informative signals, we defined a broader class of “predictive genes.” These genes are specific to an individual and a time point, and they correctly rank that individual when the ordering inferred from the other two time points is used as a reference. Unlike perfect predictors, they do not require the remaining siblings to maintain a consistent order.

Using this definition, each litter contained several thousand predictive genes. The numbers ranged from 5,983 in litter 16-30 to 7,380 in litter 16-90 (Fig. 2G). On average, about 1,000 predictive genes were shared across different time points within a given individual, whereas perfect predictors were far fewer and required full consistency across all time points. For illustration, the predictive genes for sibling β of the 16-90 litter at time point t3 showed largely preserved rank ordering at the earlier time points t1 and t2, particularly for genes at the top or bottom of the ranking (Fig. 2H).

Overall, these results show that identity signatures are not confined to the small set of perfect predictors. Instead, thousands of genes contribute to individual-specific ranked expression patterns, with a substantial fraction showing consistency across time within the same sibling.

### Individual Identity Arises from Modest Shifts in a Small Co-expressed Gene Set

To dissect the structure of identity signatures, we asked whether individuality emerges from a handful highly informative “hub” genes or from the cumulative effect of many modestly varying genes. To test this, we performed two complementary simulations based on our empirical model. First, we progressively spiked the expression data with increasing numbers of perfectly predictive genes and calculated the resulting predictability score (Fig. 3A). The empirical predictability score observed in the real data (mean = 1.8) was achieved when incorporating only 80-190 predictive genes. Relative to a rank-order null model, armadillo quadruplets contained an excess of approximately 150 predictive genes (Fig. 3B). In the second simulation, we systematically varied two parameters - the number of affected genes (range: 10 to 1,000) and their absolute effect size (|log₂FC| = 0.1 to 0.5) - and assessed the resulting predictability (Fig. 3C). The empirical score of 1.8 was reached at 150 genes, each with a modest effect size of about |log₂FC| ≈ 0.3. Thus, subtle yet coordinated expression shifts across a relatively small subset of about 150 genes appear sufficient to encode individual identity.

**Fig. 3.**
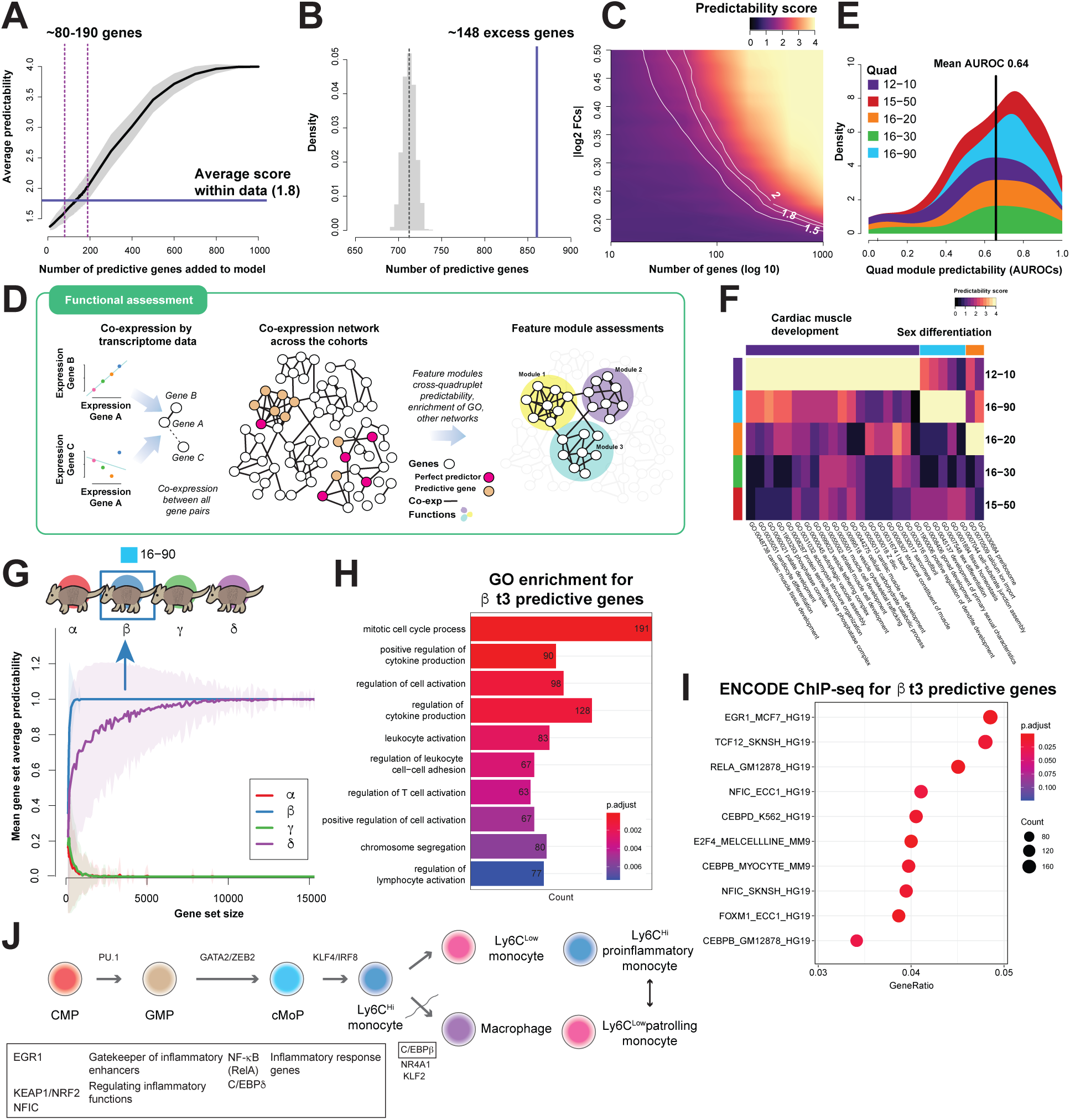
Functional characterization of identity-predictive gene modules. (**A**) A model and (**B**) our empirical estimates suggest ∼150 excess genes would contribute to the identity predictable signal. (**C**) This reflects ∼0.3 |log2FC| according to our model. (**D**) Feature genes mark identity and form feature modules (co-expressed). (**E**) Module predictability signal has weak cross-quad predictability. (**F**) These identity genes contribute to known pathways and processes that are also quadruplet specific. **G**) The identity predictability comparison among 16-90 highlights the distinctive predictability of individual *β*. Enrichment analysis of 16-90 *β* predictive genes at t_3_ for (**H**) GO annotations and (**I**) ENCODE ChIP-seq binding targets for human. (**J**) TFs enriched for 16- 90 predictive genes are associated with the differentiation process of monocytes.

To investigate whether identity signatures were associated with distinctive gene-gene relationships, we constructed gene co-expression networks for each cohort and time point separately, resulting in a total of 15 networks, which were subsequently aggregated into a single network (Fig. 3D). To assess whether identity-predictive genes exhibited stronger co-expression patterns, we applied EGAD (Extending ‘Guilt-by-Association’ by Degree), a method designed to evaluate the performance of guilt-by-association predictions within gene networks (*13*). Cross-quadruplet testing revealed that the sets of perfect predictor genes were only modestly predicted by these co-expression relationships (average AUROC = 0.64, Fig. 3E), suggesting limited consistency of interactions among these genes across different quadruplets. Additionally, analysis of expression levels indicated that perfect predictor genes were not distinct in their average expression compared to other genes (Fig. S2).

Functionally, pathway enrichment analysis identified several pathways related to cardiac muscle development and sex differentiation as differentially regulated in specific quadruplet sets (Fig. 3F), whereas their co-expression relationships are conserved between human and armadillo (Fig. S5), highlighting unique biological processes potentially underlying identity-associated variation.

### Immunological Phenotypes Linked to Predictable Individual Identity in 16-90

Identity-predictive gene sets overlapped little between the five cohorts, implying that individuality signatures are largely litter-specific. To pinpoint the origins of one such signature, we focused on quadruplet 16-90, which was phenotypically unusual: it exhibited the highest CV across haematological measures and had more perfect predictor genes than expected. Hereafter, we denote the four siblings of quadruplet 16-90 as α, β, γ and δ.

Rather than averaging predictability across the litter, we assessed each sibling separately. By evaluating the individual transcriptomic predictability rather than quadruplet predictability, individuals β and δ showed the greatest predictability (Fig. 3G), with β emerging as easily identifiable, requiring only a small number of genes. Concordantly, β displayed elevated platelet and white-blood-cell counts, consistent with an activated immune profile among the quadruplets (Fig. S6). Predictive genes for β at t_3_ were significantly enriched for GO terms related to cytokine production and to leukocyte and lymphocyte activation (Fig. 3H; Fig. S7), linking β’s transcriptomic distinctiveness to an immune-biased physiological state.

To uncover the upstream drivers of this immune-skewed signature, we next examined whether the β-specific identity genes share common regulatory inputs. Promoter analysis revealed significant enrichment of 18 transcription-factor (TF) target sets (q < 0.05; Fig. 3I, Fig. S8), with the strongest signals arising from factors that guide blood-cell differentiation - particularly within the monocyte lineage. Across individuals, the most consistently enriched targets belonged to TFs that regulate myeloid development. Foremost among them were targets of EGR1, a master regulator of inflammatory macrophage enhancers and monocyte maturation, implying a central role for EGR1 in establishing divergent expression programs (Fig. 3K and J) (*14–16*). Additional enriched TFs included RELA (NF-κB), CEBPδ, NFIC, and FOXM1, each with well-documented functions in immune and inflammatory pathways (*17–19*). Together, these findings suggest that individuality signatures are orchestrated by specific regulatory networks, particularly those governing innate immune responses.

Combined, our bulk-transcriptomic analyses show that genetically identical armadillos acquire stable, individual-specific expression profiles that persist over time and withstand moderate environmental variation. These individuality signatures are encoded by largely non-overlapping gene sets and can be explained by differences in regulatory networks rather than technical noise. The most striking example is sibling β of the 16-90 litter, whose transcriptomic distinctiveness coincides with elevated white-blood-cell counts observed in the hematological tests, hinting at an underlying shift in immune-cell composition. To investigate this hypothesis at higher resolution, we next employed single-cell sequencing analyses.

### Single-Cell Profiling Links Individuality to Immune Cell Composition Differences

To probe the cellular basis of individuality, we collected peripheral blood mononuclear cells from the 16-90 quadruplets, selected for its pronounced bulk-level variability, at a later time point (t_4_; Fig. 1F). After stringent filtering, we obtained high-quality profiles for 16,093 single cells by scRNA-seq and 19,003 nuclei by scATAC-seq. The scRNA-seq cells were almost evenly distributed across the four siblings (Fig. 4A; Fig. S9, S10), whereas the scATAC-seq dataset was skewed, with sibling β contributing a disproportionate share of nuclei both before and after filtering (Fig. S11, S12).

**Fig. 4.**
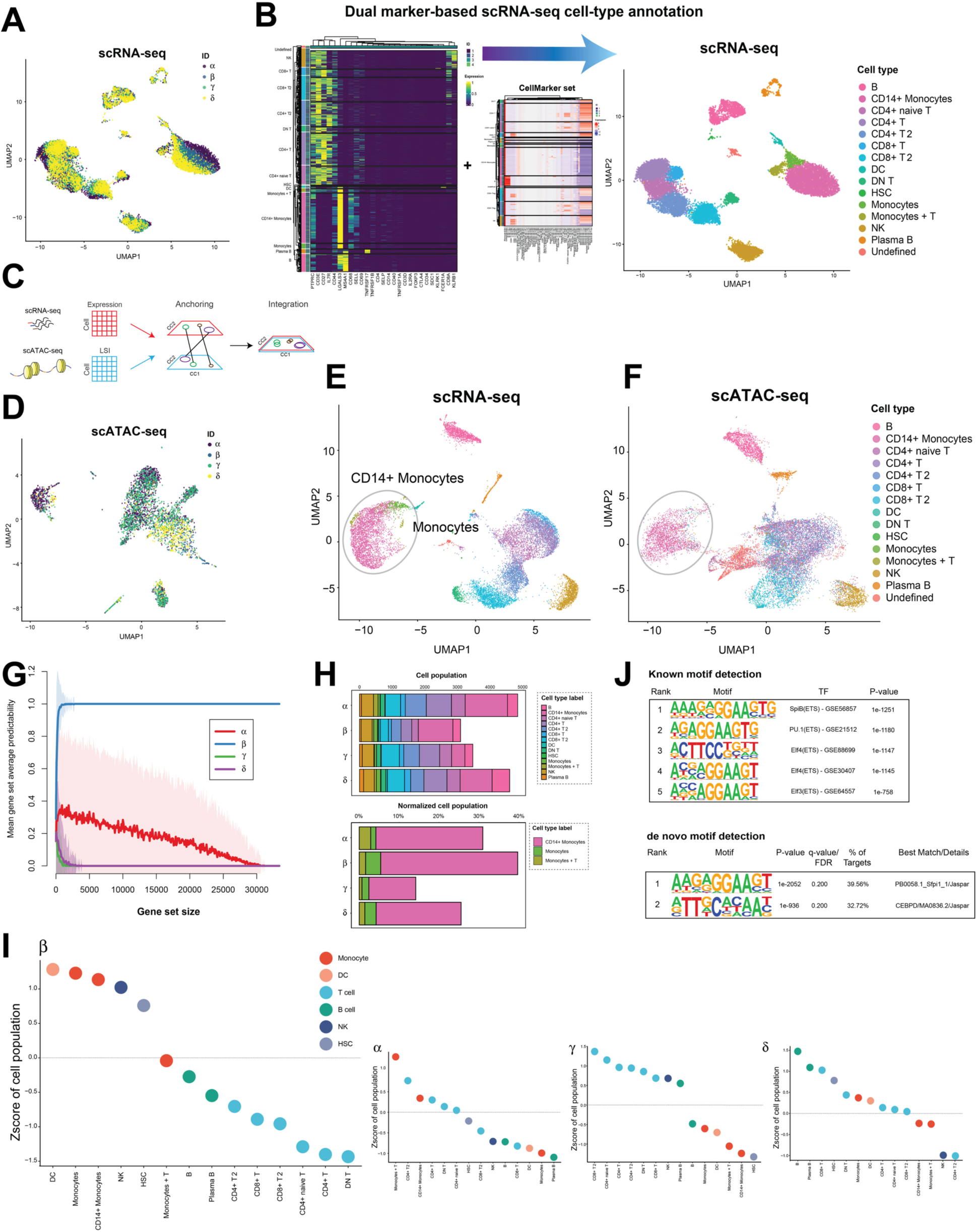
Single-cell transcriptome and epigenome analysis reveals consistent leukocyte population changes. (A) UMAP embedding of scRNA-seq data from the 16-90 lineage. Each point represents a cell, colored by individual. (B) Each cluster from scRNA-seq data is annotated according to the expression of known marker genes and module activity of CellMarker gene sets. (C) Integration of scRNA-seq and scATAC-seq was performed using Seurat. (D) UMAP embedding of scATAC-seq data. (E,F) UMAP visualization of scRNA-seq (E) and scATAC-seq gene activity profiles (F) in the integrated space with the predicted cell type annotation transferred from scRNA-seq. (G) The identity predictability comparison among 16-90 based on scRNA-seq pseudo-bulk profile demonstrates the predictability of β. (H) Horizontal stacked bars showing the distribution of cell-type counts for each individual (top) and normalized monocyte-related cell-type proportions (bottom). (I) Deviation of each cell-type population against global population for β and the other three individuals. (J) Motif analysis of CD14+ monocyte cluster using HOMER.

To identify cell types, we leveraged known marker genes and a curated database of cell type signatures. No established armadillo-specific markers for blood cell types exist, and marker gene expression can vary across species. We therefore applied a dual annotation strategy: we manually selected a panel of 27 canonical marker genes for mammalian blood cell types, and we also consulted the CellMarker database of human/mouse cell type markers (*20*). By examining the expression of these markers, we successfully annotated 14 of the 15 transcriptional clusters as known blood cell types (Fig. 4B; one small cluster remained ambiguous). The use of cross-species marker data increased our confidence in these annotations despite species differences.

We next integrated the scRNA-seq and scATAC-seq datasets to corroborate the cell type identities and investigate chromatin differences. Using Seurat-based integration, we co-embedded the transcriptomic and chromatin profiles in a common space (Fig. 4C-F). In this integrated visualization, cells grouped by cell type rather than by modality, indicating that the ATAC profiles of, say, monocytes aligned well with the RNA profiles of monocytes, etc. This allowed us to transfer labels and confirm that the scATAC-seq clusters corresponded to the same cell types identified in scRNA-seq.

To directly link the single-cell findings back to the bulk transcriptome signatures, we created “pseudo-bulk” profiles for each individual by aggregating their single-cell transcriptomes. We then asked whether the identity predictability from the bulk analysis was consistent in the pseudo-bulk profiles. Indeed, applying our supervised learning approach to the pseudo-bulk profiles successfully reproduced the strong predictability of individual β (Fig. 4G), whereas the other individuals were less distinctly classified. The integrated analysis also revealed that the overall cell type composition captured by scATAC-seq was similar, with one caveat: we noticed that the raw cell yields in scATAC-seq showed a slight bias (before filtering, one sibling had fewer high-quality nuclei). After filtering, however, the distribution of cells by individual was roughly even in scRNA-seq and only modestly uneven in scATAC-seq (Fig. S10, S11), so any composition differences are likely biological rather than technical.

With cell type labels in hand for each cell, we calculated the Z-score of each major cell type proportion for each sibling. The results were striking: the four siblings showed noticeable differences in the makeup of their blood immune cells (Fig. 4H). In particular, sibling β (the transcriptomically distinctive individual) had a substantially higher fraction of monocytes compared to the others (Fig. 4I). Monocyte-lineage cells (including classical CD14⁺ monocytes and a cluster of cells we labeled “Monocytes+T”) were overrepresented in β. This pattern is consistent with the blood hematology results including more white blood cells at t2 and t3 (Fig. S13). Correspondingly, β had slightly lower percentages of lymphocytes (T and B cells) than some of its siblings. Notably, sibling δ (the second-most predictable individual in bulk expression) showed a mild increase in T cells relative to the others, and sibling γ showed a mild increase in B cells. These shifts were smaller than the monocyte expansion in β but followed a pattern: each sibling appeared to have a unique immune cell type profile, with β’s deviation in the myeloid compartment being the most pronounced. Thus, the single-cell data confirm that functional differences in blood cell composition exist among the siblings and contribute to their transcriptomic individuality.

### Single-Cell Chromatin Accessibility Profiles Support Cell Typing Through TF Motif Enrichment Analysis

We further analyzed the scATAC-seq chromatin landscapes to identify TF motifs enriched within each cell cluster. Due to sampling bias and data sparsity, several populations detected in scRNA-seq - DC, HSC, and the mixed monocyte + T cluster - were absent from the scATAC-seq data. While DN T, Monocytes, and Plasma B clusters were captured in numbers too small for motif analysis. Focusing on clusters annotated as CD14+ monocytes, we found significant enrichment of PU.1 (encoded by SPI1) and Spi-B motifs in differentially accessible regions (DAR); these factors bind to highly similar motifs (Fig. 4J). ELF-family motifs rank within the top five enrichments for CD14+ monocytes—a contrast to human datasets where these motifs are usually more accessible in B cells than in monocytes - an effect we do not observe (Fig. S14). Collectively, the chromatin signatures reinforce our monocyte annotation across the matched scRNA-seq and scATAC-seq profiles.

### Potential Role of Cell-Type Compositional Changes in Driving Transcriptomic Identity Signatures

To trace the bulk-level identity signatures to their cellular origins, we compared bulk RNA-seq signatures with the single-cell data from the 16-90 litter. Aggregate expression of β’s predictive genes at t_3_ and other time points is markedly enriched in monocyte clusters (Fig. 5A; Fig. S15). Intersecting the 16-90 sibling-specific identity genes with the full set of predictive genes identified across all litters and time points (bulk and pseudo-bulk) again yields the strongest overlap for β (Fig. 5B). A parallel comparison with curated cell-type markers shows that β’s predictors intersect most with monocyte-specific genes (Fig. 5C).

**Fig. 5.**
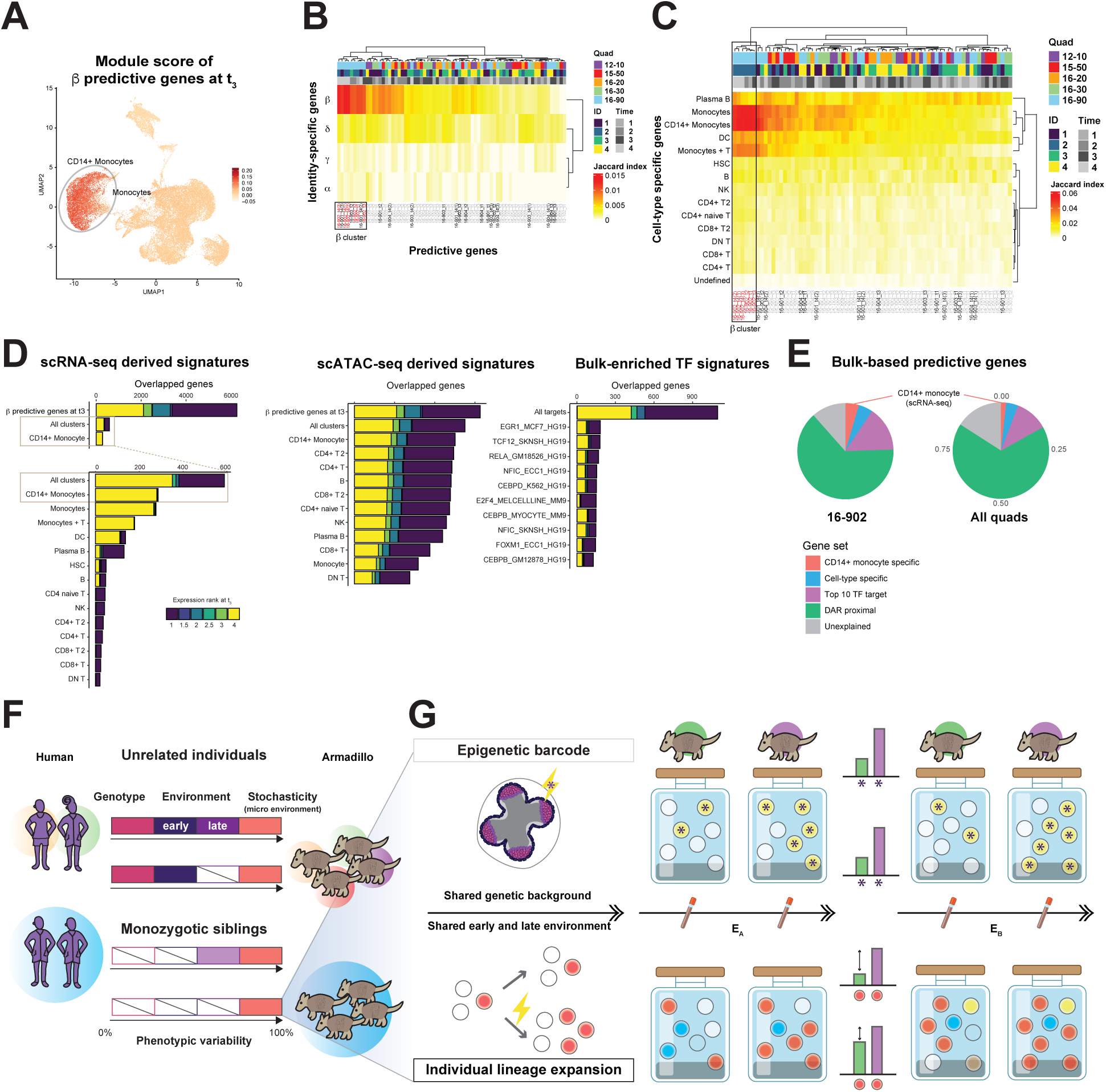
Cross-modality categorization of predictable genes indicates overlap of monocyte-derived signatures and β predictive genes. (A) UMAP of scRNA-seq data at t_3_ showing module scores for β predictive genes. (B,C) Overlap between the predictive gene sets and sibling (B) and cluster-specific (C) gene sets shows the enriched signal in β and monocyte-related clusters. The sample from 16-90 β, any other 16-90 sibling, and unrelated individual is shown in red, black, and gray, respectively. (D) The ranked expression profiles of overlapping genes with β predictive genes at t_3_ for the signatures derived from scRNA-seq (left), scATAC-seq (center) cluster-specific genes and top 10 bulk-enriched TF target genes (right). (E) Proportion of β predictive genes overlapping with: CD14+ monocyte-specific genes, other cell-type specific genes in scRNA-seq data, top 10 TF target genes, differentially accessible region (DAR) proximal genes in scATAC-seq data, and none of the above (labelled as ‘Unexplained’). (F) Phenotypic variability arises from three major sources: genetic variation, environmental variation (both early and late), and stochasticity. (G) Two potential mechanisms to promote stochastic developmental canalization: an epigenetic fingerprint established during early development, and lineage expansion occurring in later stages.

To further characterize the identity signature of sibling β, we examined the distribution of expression ranks for its predictive genes at t_3_. The monocyte-specific genes tended to occupy the extremes—either highest (rank 4) or lowest (rank 1)—relative to the other siblings (Fig. 5D). The number of those genes surpassed the number of theoretically estimated identity signatures for observed bulk data, specifically 80-190 or 150 genes by two different simulations, suggesting the monocyte enrichment possesses a sufficient power to make sibling β predictable in bulk. Notably, the subset of monocyte-specific genes within this predictive set was predominantly highly expressed in β, reinforcing the cell-type–biased pattern.

In contrast, genes located near differentially accessible regions (DAR-proximal genes) showed more modest differences across cell types, with a rank distribution similar to that of the full β predictive set (Fig. 5D). Although the sizes of these gene sets vary, this trend suggests that peak-level signals from scATAC-seq may not always reflect distinct cell-type–specific regulation, possibly due to redundancy in regulatory elements. Additionally, the targets of the top 10 TFs identified from the bulk data were enriched for genes with extreme expression ranks with a bias toward lower-ranked genes.

To integrate these findings, we organized identity-predictive genes for β and all individuals into four mutually exclusive categories, applied in this order: CD14⁺ monocyte-specific genes, genes specific to other cell types, top 10 TF targets, and DAR-proximal genes (Fig. 5E). Among β’s predictive genes, 4.6% were classified as CD14⁺ monocyte-specific, compared to just 1.7% across all cohorts, further emphasizing the monocyte-skewed nature of β’s transcriptomic identity. Together, these findings indicate that the transcriptomic signature distinguishing sibling β is driven primarily by a monocyte-associated gene program.

## Discussion

Our results show that transcriptomic individuality in genetically identical animals corresponds to stable differences in immune-cell composition. In nine-banded armadillo quadruplets raised under controlled conditions, each sibling developed a distinct gene expression profile that persisted over time. These profiles were not limited to neutral molecular markers. Instead, they reflected coordinated differences in cell-type abundance that were evident at both bulk and single-cell levels. The most striking case was sibling β of the 16-90 litter, which displayed a consistent expansion of monocyte-lineage cells and an associated innate immune program. This establishes that non-genetic individuality can produce durable physiological divergence in immune composition.

Our earlier work identified allele-specific expression (ASE) imbalance as a stable molecular fingerprint of individuality in the same animals (*10*). That signal arose from persistent monoallelic biases and did not require changes in total gene output. In contrast, the individuality described here is driven by differences in total expression levels. We found little overlap between ASE-based and total-expression-based signatures, which suggests that they represent independent axes of non-genetic variation. ASE variation likely captures neutral epigenetic differences, while the expression signatures identified here reflect functional shifts that influence cell physiology and abundance.

The nature of these shifts offers insight into possible mechanisms. The single-cell data point to subtle differences in lineage production or survival rather than dramatic reprogramming. Modest changes affecting about 150 genes were sufficient to recapitulate the observed identity signal in simulation, and these genes exhibited coordinated but small expression shifts. These patterns are compatible with the idea that early stochastic events in hematopoietic development bias the relative representation of particular lineages. Such processes need not involve somatic mutations. Reanalysis of ASE data found no evidence for reproducible allelic imbalance at canonical clonal hematopoiesis genes, and the animals were much younger than the age at which clonal expansions typically arise in species with comparable lifespans. Although somatic mosaicism cannot be fully excluded, the available evidence suggests that the individuality observed here derives mainly from developmental variation rather than mutational clonality.

The broader implications of these findings relate to non-genetic individuality more generally. Stable biases in immune-cell composition have been documented among monozygotic human twins, and these differences are often attributed to non-heritable influences early in life (*1*, *21*). Our results provide a model in which such variation can arise without genetic or major environmental differences (Fig. 5F and G). Early stochastic cues during development can produce long-lasting shifts in lineage balance, which then manifest as stable differences in transcriptomes and immune profiles. This framework provides a potential explanation for why genetically identical individuals sometimes diverge in disease susceptibility, including autoimmune and inflammatory conditions. More speculatively, developmental sources of heterogeneity among genetically identical individuals could serve as a form of bet hedging, increasing the resilience of a population to diverse pathogens (*22*). Although our study was not designed to test adaptive hypotheses, the presence of individualized immune profiles in a clonal sibling group demonstrates that developmental stochasticity can generate phenotypic diversity with potential functional consequences.

In summary, genetically identical armadillos raised in nearly identical environments acquire distinct and stable transcriptomic signatures that reflect coherent differences in immune-cell composition. These differences are consistent over time and are detectable at both bulk and single-cell resolution. They arise independently of ASE variation and do not require somatic mutations in known clonal expansion pathways. Together, these findings illustrate how early stochastic influences can become embedded in the transcriptomic and cellular landscape, giving rise to lasting and biologically meaningful individuality.

## Materials and Methods

### Armadillo Collection and Sample Acquisition

Five sets of nine-banded armadillo (*Dasypus novemcinctus*) quadruplets (20 animals total) were used in this study. These animals were obtained and reared under controlled conditions as described previously (*10*). Briefly, pregnant females were captured from the wild and immediately transported to the holding facility at the University of the Ozarks, Clarksville, Arkansas. The pups were kept with the mothers until about 6-10 weeks postnatal age, where most litters shared the pen with another litter (2 litters per pen). After separation from the mothers, at four to five months of age, the animals were delivered to the National Hansen’s Disease Program (NHDP) facility in Baton Rouge, LA, where they were placed in pairs in modified rabbit cages.

Blood samples for transcriptome analysis were collected at three time points per quadruplet staggered over the course of a year (Fig. 1F), starting from March 2017 until August 2018. The pilot study was performed for two sets of quadruplets (12-10 and 15-50), and then later quadruplets 16-20, 16-30 and 16-90 were included into the subjects. Only for 16-90, the blood samples were additionally extracted for single-cell analyses in Aug 2019. The original two of the five sets of armadillo quadruplets were infected intravenously in the saphenous vein with 1×10^9^ *Mycobacterium leprae* after the first time point. The animals were humanely sacrificed when they developed heavy *M. leprae* dissemination with severe hypochromic microcytic anemia. Otherwise, the animals will eventually succumb to secondary complications of persistent bacteremia because of the bacteria located in the bone marrow.

### Human twin samples

Transcriptomic data from blood samples in the TwinsUK cohort were used for the human twin analysis (*23*). Briefly, BAM files were obtained from the EUROBATS project (EGAD00001001088) following approval for access to controlled data. The dataset includes 391 blood samples collected from monozygotic (MZ) and dizygotic (DZ) female twin pairs. After quality control, 66 MZ and 96 DZ twin pairs were used for our analysis.

### Armadillo DNA- and RNA-sequencing data

We used RNA-seq data generated in our previous study (*10*), which is publicly available through the Gene Expression Omnibus (GEO) under accession number GSE141951, with further details on read processing and genome construction provided in that study. For quantification, RNA-seq reads were aligned to personalized genomes constructed from matched DNA-seq data using STAR. Transcript abundance was normalized as counts per million (CPM) for comparison across samples.

### Armadillo single-cell RNA-seq and single-cell ATAC-seq

For each sample, single-cell RNA-seq (scRNA-seq) libraries were prepared with the 10x Genomics Chromium Single Cell 3′ Gene Expression (Next GEM) kit, targeting ∼20,000 cells per sample. Single-cell ATAC-seq (scATAC-seq) libraries were generated with the 10x Genomics Chromium Single Cell ATAC Library & Gel Bead Kit (Next GEM). Multiplexed paired-end sequencing (PE76) was carried out on an Illumina NextSeq 500 on multiple flow cells.

Demultiplexing and barcode processing were performed using Cell Ranger (v3.0.2, 10x Genomics), with each sample processed individually. Reads were mapped to the armadillo genome (DasNov3, Ensembl release v95), and genome indices were generated using STAR (v2.7) (*24*). For scRNA-seq analysis, we used STARsolo with the FASTQ files as input to generate gene-count matrices (*25*). Following careful inspection of the cell-typing results, we decided to replace the Ensembl gene definition of CD4 (JH562564:228,438–255,355) with the RefSeq annotation (JH562564:226,782– 265,885). This decision was based on the weak signal observed in our annotated scRNA-seq data with the Ensembl definition, likely due to a >1,600 nucleotide difference between the transcription start sites (TSS) of the two annotations. Raw sequencing data and processed object files of scRNA-seq and scATAC-seq analysis are publicly available at the GEO under accession number GSE273197.

### Single-cell RNA-seq data analysis

Gene count matrices from PBMC samples of the 16-90 siblings were processed using the Scanpy toolkit (*26*). After preprocessing, 16,093 cells were retained for downstream analysis and assigned to 15 clusters using Leiden clustering (*27*). Detailed preprocessing steps are provided in the accompanying Jupyter notebook available in our GitHub repository. Summary statistics and basic properties of the scRNA-seq data are presented in Fig. S11 and S12.

### Single-cell ATAC-seq analysis

Raw BAM files were imported and converted into Snap objects using SnapATAC (v1.0.0) (*28*). From the mapped reads, SnapATAC generated bin-by-cell, peak-by-cell, and gene-by-cell matrices. For the bin-by-cell matrix, a bin size was set to 5 kb. Peak-by-cell matrices were constructed using peak calls from MACS2. To generate gene-by-cell matrices, the 1 kb bin-by-cell matrix was first binarized, and counts were summed for each Ensembl gene (and for CD4, the RefSeq annotation was used) by aggregating bins located within the gene body and promoter region (defined as 2,000 bp upstream of the transcription start site).

To remove low-quality cells, we applied a filtering threshold of ≥1,500 reads per cell and a UQ score ≥2.35, representing the number of unique molecular identifiers (UMIs) in the mapped reads. After filtering, 19,003 cells were retained for downstream integration with the scRNA-seq data. Normalization, clustering, and UMAP visualization were performed with the SnapATAC software package. Summary statistics and quality metrics of the scATAC-seq data are shown in Fig. S11 and S12.

### Cell-type annotation

To assign cell-type identities to the 15 scRNA-seq clusters, we first computed a pseudo-bulk expression profile for each cluster and visualized expression patterns of manually curated marker genes (Fig. 4B). After aligning gene synonyms and excluding genes with low or missing expression, 27 marker genes were used to identify blood cell types: PTPRC, CD3D, CD3E, IL7R, IL2RA, CD4, FOXP3, CTLA4, CD8A, CD27, MS4A1, TNFRSF17, SDC1, FCER1A, CD40, KLRB1, KLRK1, CD34, SELL, CD14, SELP, LGALS3, CD68, CD44, CD69, TNFRSF1A, and TNFRSF1B. As a complementary approach, we retrieved marker gene sets for human and mouse blood cells from the CellMarker database (*20*). To assess the enrichment of these gene sets among upregulated genes in each cluster, we computed module scores using the AddModuleScore function in Seurat v4 (*29*) (Fig. 4B, middle panel).

### Single-cell integrative analysis and label transfer

To jointly embed the scRNA-seq and scATAC-seq datasets, we performed canonical correlation analysis (CCA)-based integration using Seurat (*29*). For the scATAC-seq data, gene-level chromatin activity was represented by the gene-by-cell matrix generated by SnapATAC, which was incorporated as a feature in the corresponding Seurat object. Cell-type labels were then transferred from the annotated scRNA-seq data to the scATAC-seq cells using Seurat’s FindTransferAnchors and TransferData functions.

### Measuring transcriptional stochasticity

To assess transcriptional stochasticity, we conducted a transcriptome-wide clustering analysis of the samples using Spearman’s rank correlation to quantify pairwise similarity. Within each quadruplet set, highly variable genes (HVGs) were identified by ranking genes based on their coefficient of variation and mean expression levels. To evaluate the recurrence of transcriptional variability across individuals, we assessed the overlap of HVGs between different quadruplet sets. For each set, we selected the top 100 HVGs at a given time point and computed their AUROC scores in cross-quadruplet comparisons. Additionally, we performed ANOVA across all samples within each quadruplet to identify genes exhibiting the highest intra-quadruplet variability (Fig. S6).

### Supervised learning to quantify the quadruplet identity predictability from expression

To evaluate the significance of transcriptomic individuality, we applied a supervised learning framework designed to evaluate the relative consistency of gene expression across time points, as described previously (*10*). Briefly, two time points were selected to form a training set, and the rest is left for testing. For each gene, a gene scoring matrix (4 by 4) is built by comparing the ordering of the test and training CPM data averaged for two samples. Each individual gives a score of 1 to the test data individual whose rank it matched, and 0 otherwise. Then all the matrices for the selected genes feature are aggregated to produce one scoring matrix, where we associate each individual in the test data with that in training according to the scores following a winner-takes-all strategy. We repeat this three times for the different combinations of the training and test data. The final score of “predictability score” is between 0 and 4, with 4 as perfect predictability i.e., each armadillo correctly identifies its future (or past) self. The feature set consists of the genes whose order is consistent across the training samples without the information of test data while a perfect predictor gene is perfectly correlated across all time points. We obtained analytic and empirical p-values for this score by convolution of the expected distributions or repeating the learning task on randomly selected genes, respectively.

### Empirical models to estimate the number of genes and lineage contributions

To investigate the underlying mechanisms driving individual identity signatures, we estimated the number of genes contributing to the observed signal using a series of empirical models (Fig. 3A–C). To generate empirical distributions of expression-based identity scores, we simulated two models. In the first model, we assumed the identity signal is localized, and introduced a perfectly predictable signal to a subset of genes in the original expression matrix. In contrast, the second model assumed the identity signal is distributed across a larger number of genes, each contributing a weaker effect. Specifically, we simulated expression-based identity predictability by adding a fixed proportion of variance (denoted M) to a defined number of genes (N). For each simulation, the average identity prediction performance was computed. The introduced signal strength (M) was converted to an absolute log2 fold change (|log2FC|) for interpretability.

### Gene co-expression network construction and functional module evaluation

For each quadruplet at each time point, we constructed a gene co-expression network (*30*) (Fig. 3D). Briefly, Spearman correlation coefficients were calculated for all gene pairs based on expression across the four individuals within each quadruplet. These correlation values were used as edge weights in a gene-gene co-expression network, which was subsequently rank-standardized. The resulting individual networks (15 in total) were summed to produce an aggregate co-expression network. For all networks, we first applied unsupervised clustering analyses and tested for functional enrichment of the network modules identified by using our supervised machine learning tool EGAD (*13*) on the aggregate network and the individual networks. In EGAD, we report performance using AUROCs, as is typical within machine learning (Fig. 3E). An AUROC is the area under the receiver operator characteristic curve, and is the probability that a positive result is ranked higher than negative result by the algorithm (e.g., does this list of DE genes rank genes involved with “function X” higher than other genes). We used the functional annotations we mapped from GO for this assessment. To test for cross-quadruplet generalizability, we employed a leave-one-out validation approach: individual networks were aggregated while excluding one quadruplet, and modules predictive of identity in the excluded set were evaluated for performance in the aggregate network using EGAD (Fig. 3E).

### Functional analysis of identity signatures in the 16-90 Cohort

Predictability scores were calculated for each individual in the 16-90 cohort and averaged across three different combinations of training and test time points (Fig. 3G). To functionally characterize genes contributing to individual identity signatures in this cohort, we performed enrichment analysis using the R interface to enrichR. Gene Ontology (GO) and ENCODE TF ChIP-seq 2015 were selected as reference databases for this analysis (*31*, *32*).

### Detecting the significance of cell-type compositional variation

After cell-type annotation of scRNA-seq, we assessed whether specific cell types were significantly enriched in specific individuals. We converted the percentage of each cell-type composition in the individual *i c_i_* into the z-score. Using the cell type composition data of all four siblings, the z-score was computed based on the equation *z_i_* = (*c_i_* − µ)/*σ*, where *μ* and *σ* is the mean and standard deviation of the distribution of *c* across four quadruplets, respectively.

### Enrichment of identity signatures across gene sets

The functional classification of the identity signatures was carried out by comparing the gene sets obtained in this study. To evaluate the overlap of two gene sets *g_i_* and *g*_j_., Jaccard index is obtained by 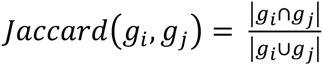 . First, we compared the identity predictive genes obtained from bulk time course data with those from scRNA-seq pseudo-bulk data. For cell-type specific genes, top 1,000 genes were selected for each cell type as the cluster-specific genes based on Wilcoxon rank-sum test p-values.

## Disclaimer

The content and views expressed in this article are solely the opinions of the authors and do not necessarily reflect the official policies of the National Institutes of Health, the U.S. Department of Health and Human Services or the Health Resources and Services Administration, nor does mention of the department or agency names imply endorsement by the U.S. Government.

## Acknowledgments

We thank Paul Pavlidis, Tony Zador, Peter Koo, Megan Crow and Dan Levy for helpful discussion and comments. The content is solely the responsibility of the authors and does not necessarily represent the official views of the National Institutes of Health. Armadillo blood for the study was obtained from the National Hansen’s Disease Program. TwinsUK is funded by the Wellcome Trust, Medical Research Council, European Union, the National Institute for Health Research (NIHR)-funded BioResource, Clinical Research Facility and Biomedical Research Centre based at Guy’s and St Thomas’ NHS Foundation Trust in partnership with King’s College London. We used ChatGPT for AI-assisted editing.

## Funding

National Institutes of Health R01LM012736 (JG)

National Institutes of Health R01MH113005 (JG)

National Institute of Allergy and Infectious Diseases Interagency Agreement IAA

15006-004 (MTP, LBA)

HRSA/NIH/NIAID/ IAA AAI25003-001-00000 (MTP)

JSPS KAKENHI Grant Number 23K14165 (RKK)

The Uehara Memorial Foundation Postdoctoral fellowship (RKK)

CiRA foundation research grant 2022 startup (RKK)

iPS academia Japan research grant 2022 (RKK)

JST ACT-X Life and Information JPMJAX24L5 (RKK)

## Author contributions

Conceptualization: RKK, SB, JG

Methodology: RKK, SB, JG

Armadillo collection: FMK

Blood collection: MTP

Investigation: RKK, SB, MTP, LF, JG

Visualization: RKK, SB

Supervision: RKK, SB, LF, JG

Writing—original draft: RKK, SB, LF, JG

Writing—review & editing: RKK, SB, MTP, LF, FMK, LBA, JG

All authors discussed, critically revised, and approved the final version of the manuscript.

## Competing interests

L.F. serves as a scientific advisor for Lighthouse Pharma, and has received consulting fees from PeopleBio Co. and GC Therapeutics Inc. The remaining authors declare no competing interests.

## Data and materials availability

RNA-seq data repurposed from our previous study (*10*) is published at GEO under accession number GSE141951. Raw sequencing data and processed object files of scRNA-seq and scATAC-seq analysis are available under accession number GSE273197. Armadillo’s trait data such as body size and blood test results were published together with the previous study (*10*). Other intermediate files can be downloaded or recreated by running the scripts found at https://github.com/carushi/ayotochtli.

## Supplementary Materials

Supplementary_data Supplementary_table 1

Supplementary_table 2

Supplementary_table 3

Supplementary_table 4

## Notes

https://github.com/carushi/ayotochtli

### Summary of Updates

Correct mislabeling in Figure 4. Correct numerical values affected by a code bug in Figure S3.

