## Supplementary materials for "Monocyte Lineage Expansion Drives Transcriptomic Individuality in Genetically Identical Armadillo Quadruplets"

Risa Karakida Kawaguchi *et al.*

**This file includes:**

Supplementary Text
Figs. S1 to S15
References (1 to 6)

**Other Supplementary Materials for this manuscript include the following:**

Data S1 to S4

**Supplementary Text**

STARsolo alignment settings

Demultiplexing and barcode processing were performed using Cell Ranger (v3.0.2, 10x Genomics), with each sample processed individually. Reads were mapped to the armadillo genome (DasNov3, Ensembl release v95), and genome indices were generated using STAR (v2.7) [1]. For scRNA-seq analysis, we used STARsolo with the FASTQ files as input to generate gene-count matrices [2]. The following options were applied: --soloType Droplet --soloCBwhitelist 3M-february-2018.txt --soloCellFilter CellRanger2.2 8000 0.99 10 --soloFeatures Gene Velocyto --quantMode GeneCounts --soloBarcodeReadLength 28. For scATAC-seq analysis, STARsolo was run in paired-end mode to generate BAM files. The options used for mapping were: --outSAMattributes NH HI CR CB --soloType CB_samTagOut --soloCBmatchWLtype 1MM --soloBarcodeReadLength 16 --twopassMode Basic --soloFeatures Gene GeneFull SJ Velocyto --soloCBwhitelist 737K-cratac-v1.txt.

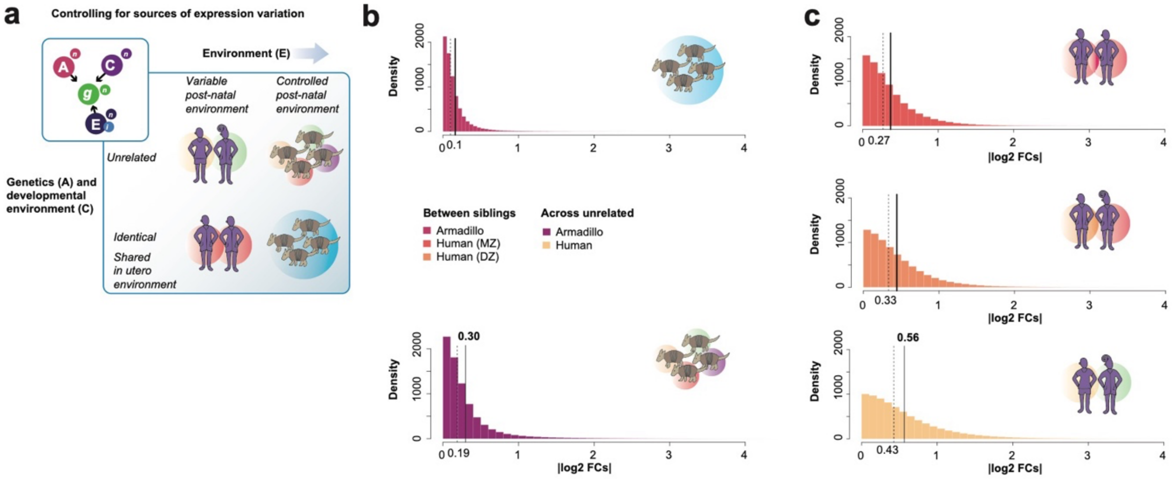


**Fig. S1.**

(A) Phenotypic variation results from three main sources: genetic variation, epigenetic variation and environmental effects. These are all influenced by stochasticity of gene expression. (B) and (C) As a measure of transcriptional (and phenotypic) variation between samples, we calculated pairwise $\log_2$ fold changes ($|\log_2 FC|$) of gene expression values. We normalized each sample to CPM, and then took the absolute value of $\log_2$ of the ratio of each gene's CPM ($g_i$ is a gene, $n$ and $m$ are individual samples). To ensure that we are comparing similar genes, we restricted our analysis to approximately 7500 genes that were homologs and were in both expression data sets. (B) Distributions of log2 fold changes calculated between armadillos within the same quadruplet, and those across quadruplets. (C) In comparison to fold changes between human twins (MZ and DZ), and those between unrelated individuals.

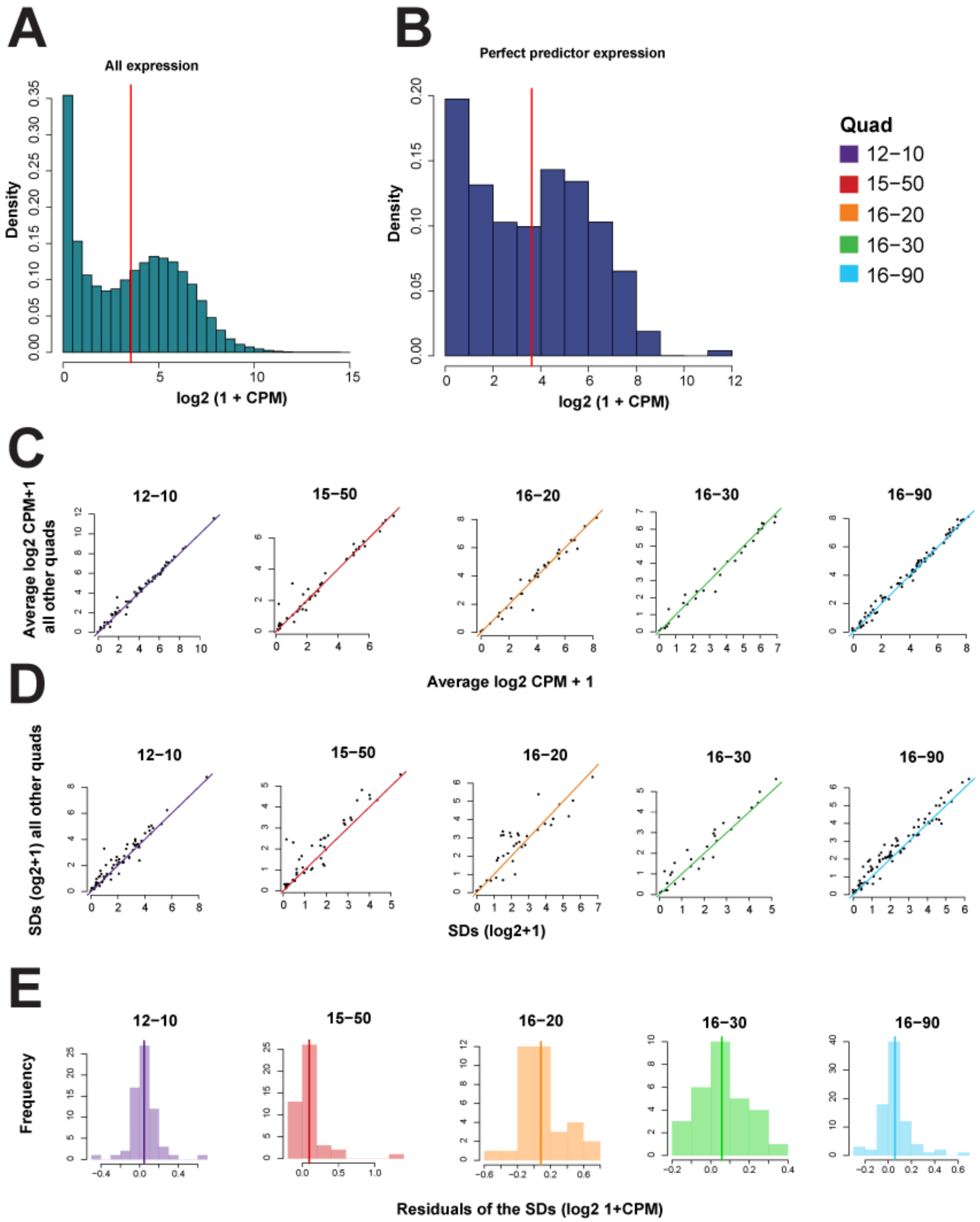


**Fig. S2.**

Characterization of perfect predictors showing normal expression properties.
(A) Expression profiles of perfect predictor genes (B) are similar to background expression as shown in (A). (C) Scatterplots of average gene expression within a quad compared to all other quads. The expression values sit very closely along the diagonal. (D) SDs of perfect predictors compared across armadillos. (E) Distributions of residuals of variance within a quad compared to all other quads. If there is excess variance, we expect to see a positive shift. On average, no signal is apparent.

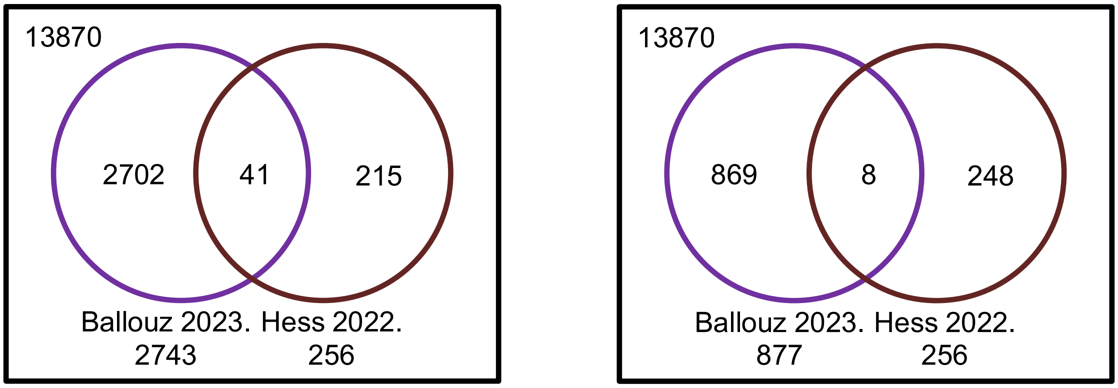


**Fig. S3.**

Venn diagram for DEGs obtained from Hess 2022 [3] and Ballouz 2023 [4]. Up-regulated genes between pre- and post-infected samples were extracted from Hess 2022 by one-way ANOVA then assigned to each blood cell type. DEGs from Ballouz 2023 were obtained by Wilcoxon signed rank test (p=0.05), and all DEGs and only up-regulated DEGs are used to draw Venn diagrams (left and right, respectively).

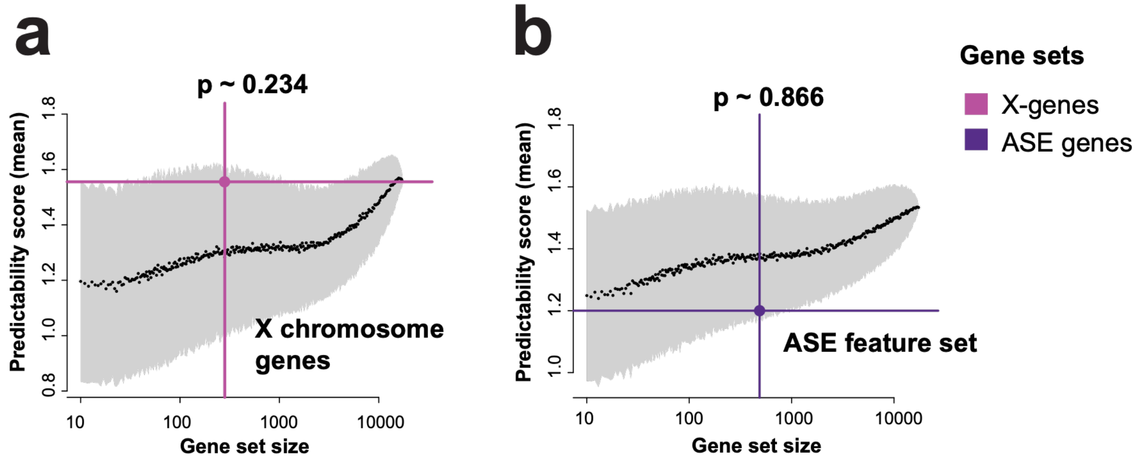


**Fig. S4.**

(A) Average identity predictability achieved by X-chromosome genes and (B) ASE imbalanced genes reported in Ballouz S, et al. 2023 [4] over the distribution of empirical null of randomly selected gene sets shown in gray.

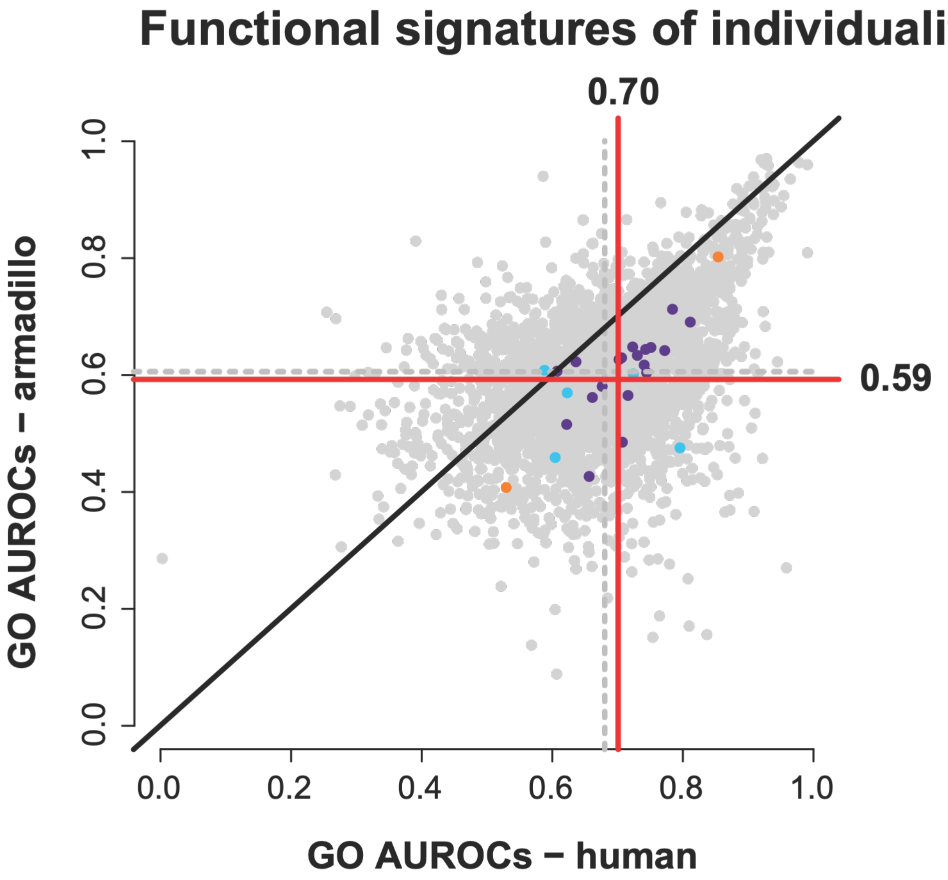


**Fig. S5.**

Co-expression relationships shared across species due to conserved stochasticity.
We constructed an aggregate human network from a large compendium of human expression data, or all extant blood data available within recount2 [5]. The human data is across 60 experiments (a total of 3,174 samples). We use "experiment" to refer to an entire expression dataset, across all its samples. After constructing the aggregate network, we used EGAD [6] to measure the network's performance with GO. We compared the AUROC GO performances across the species, including the GO gene sets enriched in the perfect predictors (points shown in the color corresponding to the cohort as in **Fig. 3F**).

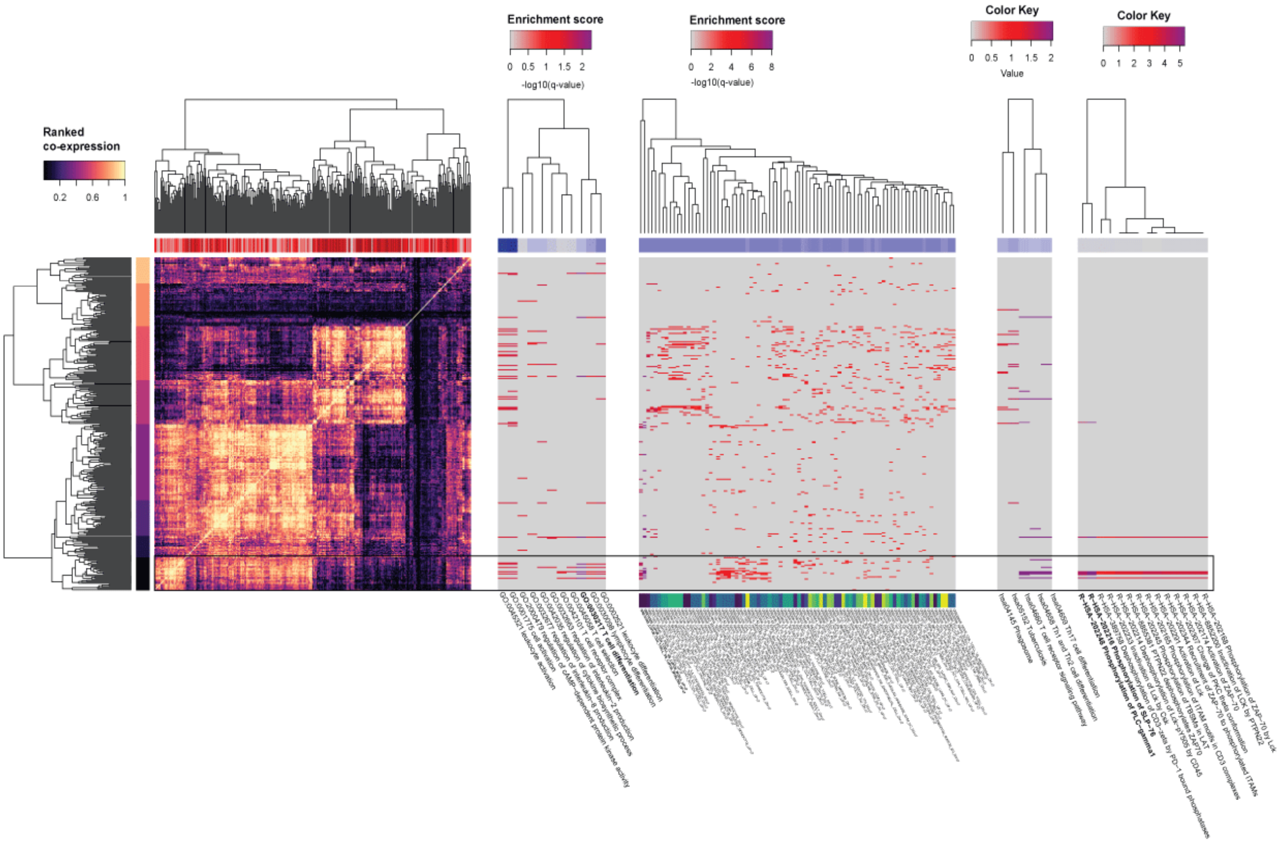


**Fig. S6.**

Gene set enrichment analysis for 16-90 DE genes. GO, MSigDB, KEGG, and Reactome pathway enrichments for the ANOVA DE genes. The strongest signal is for T-cell differentiation.

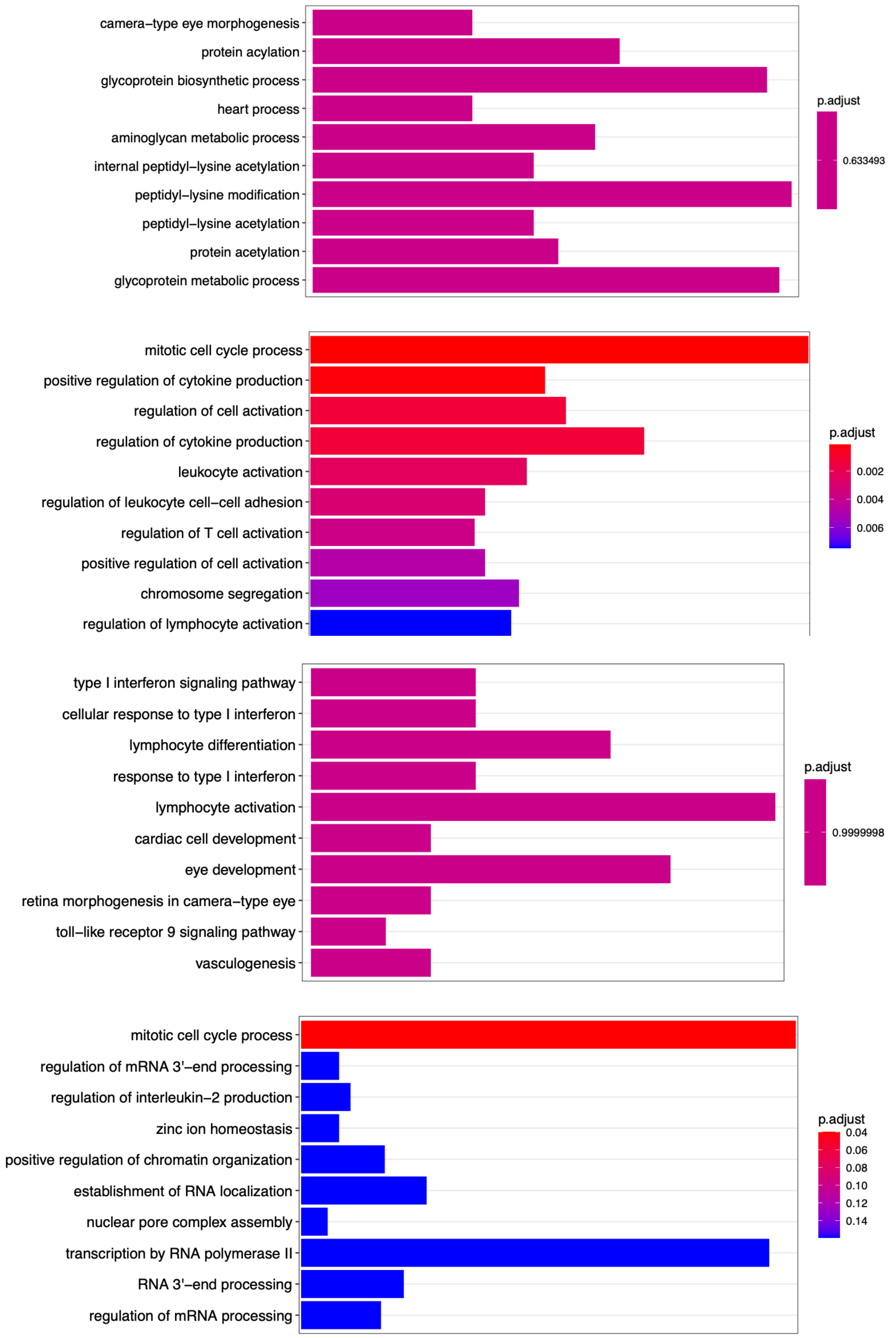


**Fig. S7.**

GO Enrichment analysis of the predictive genes at $t_3$ for 16-90 $\alpha$, $\beta$, $\gamma$, and $\delta$ from top to bottom.

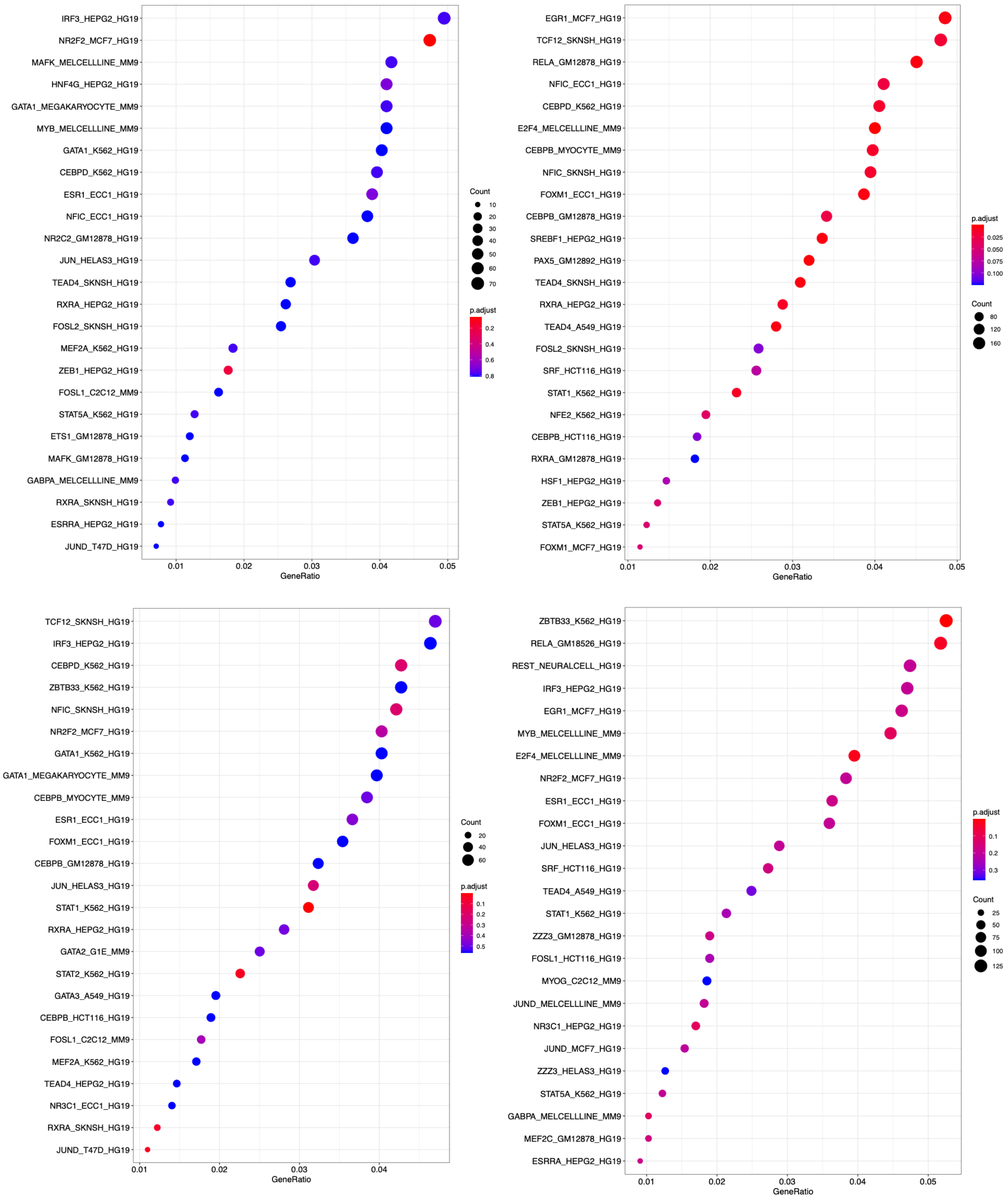


**Fig. S8.**

ENCODE human ChIP-seq binding target enrichment analysis of the predictive genes for 16-90 $\alpha$ (top-left), $\beta$ (top-right), $\gamma$ (bottom-left), and $\delta$ (bottom-right) at $t_3$.

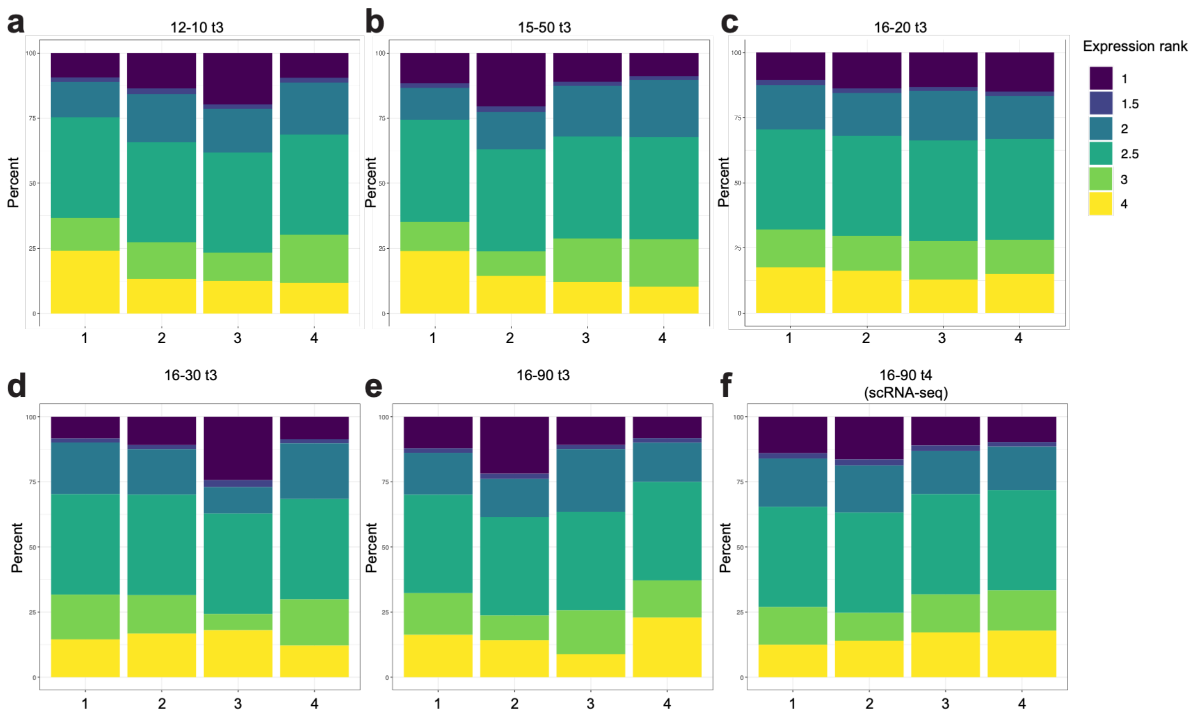


**Fig. S9.**

Expression rank distribution for each quadruplet. (A-E) Expression rank of 12-10, 15-50, 16-20, 16-30, and 16-90 at $t_3$. Mean ranks are assigned to ties. (F) Expression rank of 16-90 at $t_4$, calculated based on scRNA-seq pseudo-bulk profiles.

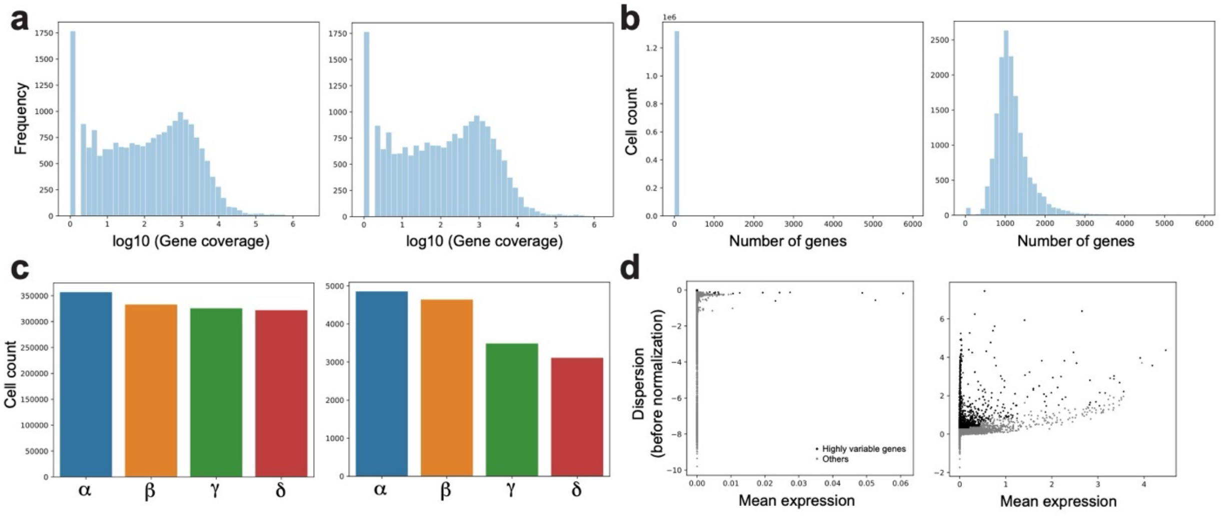


**Fig. S10.**

Basic properties of scRNA-seq data before (left) and after (right) STARsolo filtering. (A) log10 (read coverage) for each gene and (B) number of observed genes for each cell. (C) Detected cell number of each quadruplet. (D) Mean expression and dispersion of highly variable genes and others, before and after filtering.

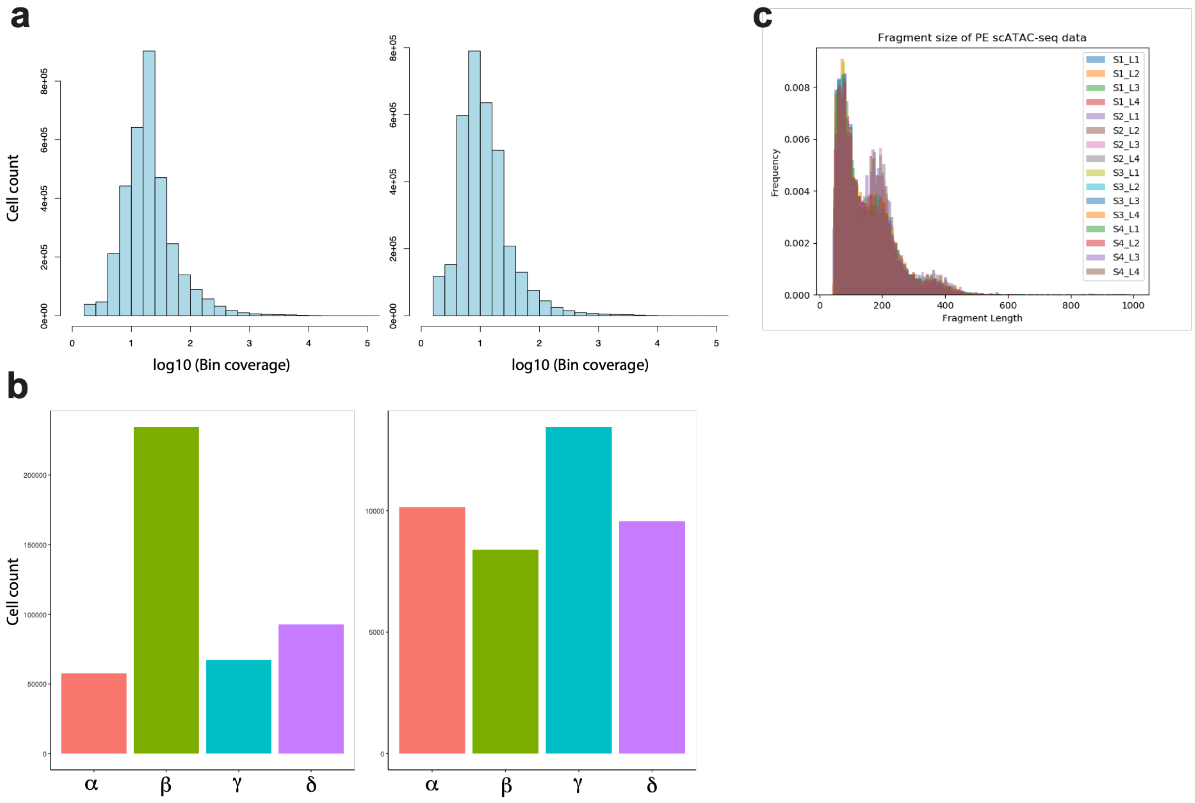


**Fig. S11.**
(A) and (B) Basic properties of scATAC-seq data before (left) and after (right) custom filtering. (A) log10 (read coverage) for each 5kb bin and (B) detected cell number of each quadruplet. (C) Fragment length distribution of paired-end scATAC-seq data from each quadruplet (S1-4) sequenced in each lane (L1-L4).

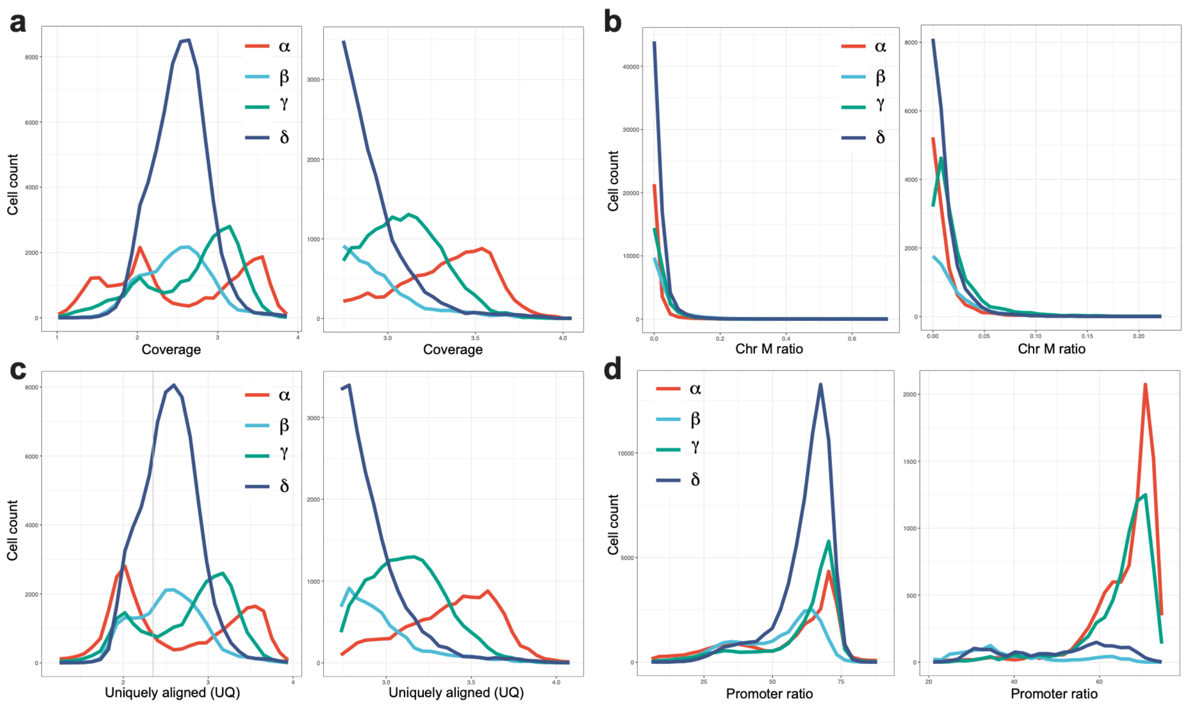


**Fig. S12.**

Basic properties of scATAC-seq data before (left) and after (right) custom filtering. (A) log10 (read coverage + 1) for each 5kb bin. (B) Ratio of reads assigned to mitochondrial chromosome. (C) log10 (read coverage + 1) of uniquely aligned fragments for each cell. (D) Ratio of fragments aligned to the promoter regions.

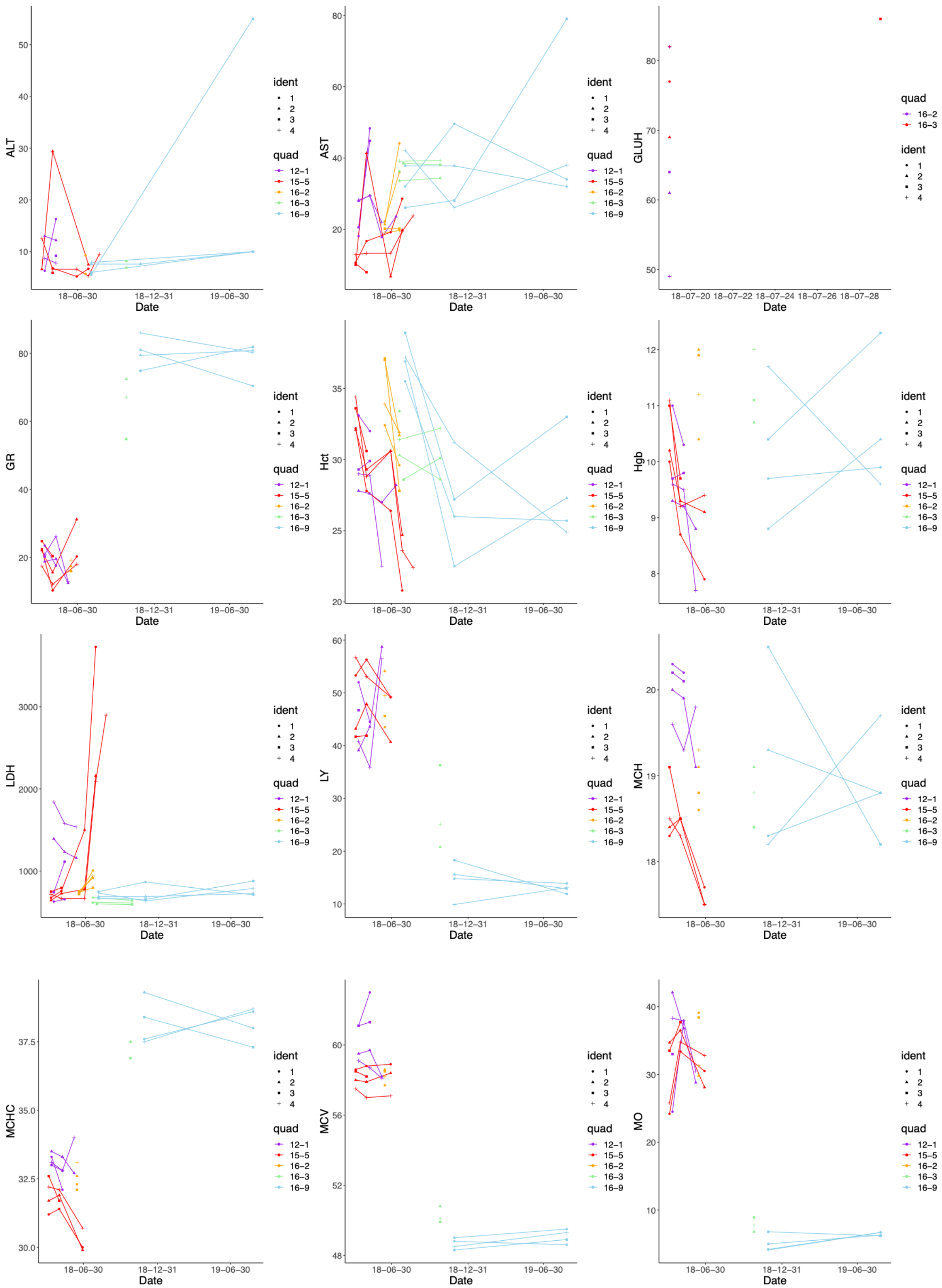
ALT
Date
18−06−30
18−12−31
19−06−30
ident
1
2
3
4
quad
12−1
15−5
16−2
16−3
16−9
AST
GLUH
18−07−20
18−07−22
18−07−24
18−07−26
18−07−28
GR
Hct
Hgb
LDH
LY
MCH
MCHC
MCV
MO

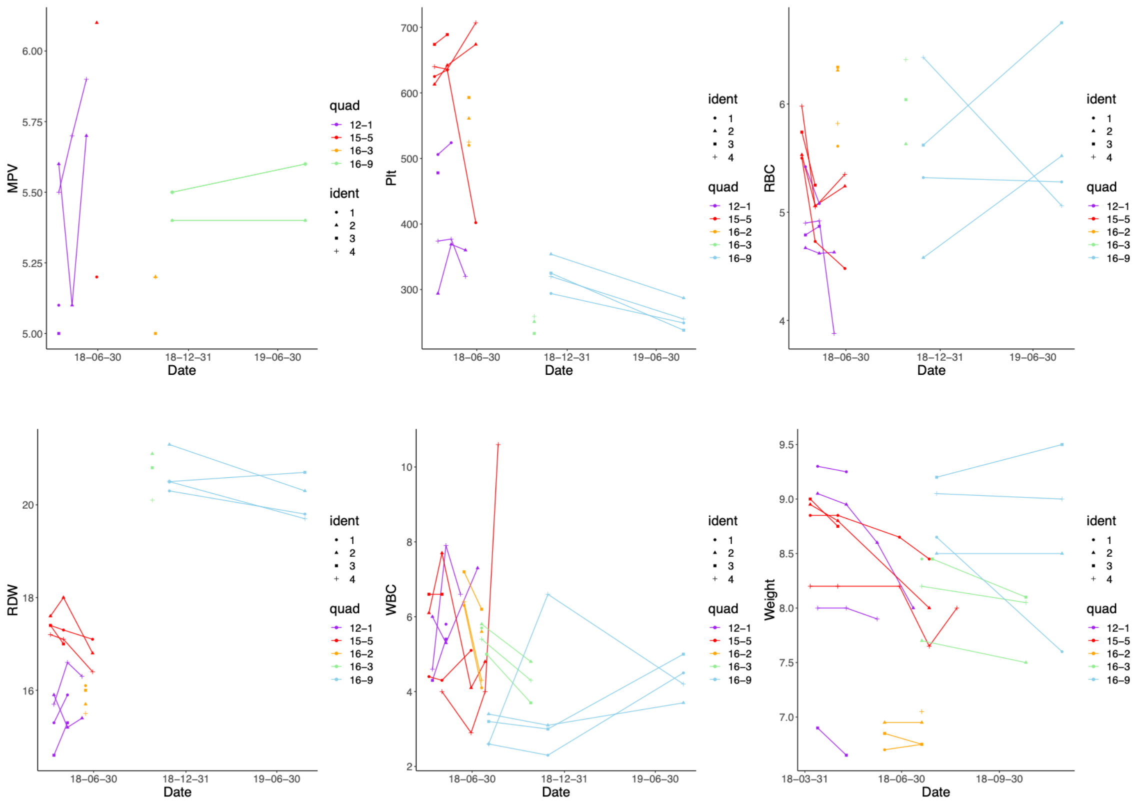


**Fig. S13.**

Time-course change of blood test and phenotypic traits of the armadillo quadruplets. ALT, AST, GLUH, GR%, HCT, HGB, LDH, LY %, MCH, MCHC, MCV, MO %, MPV, PLT, RBC, RDW, WBC, Weight.

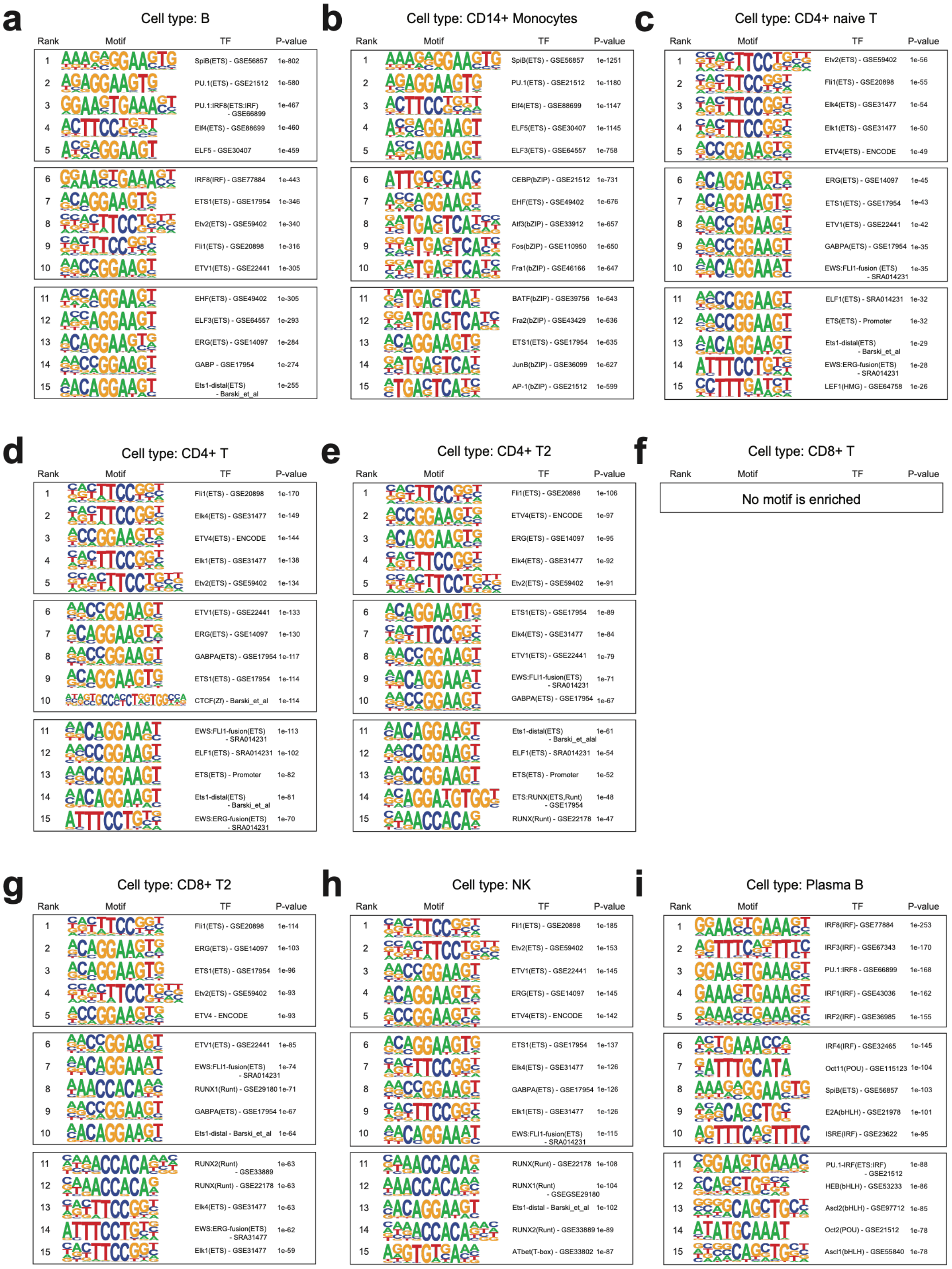


**Fig. S14.**
Detected known motifs for each cell cluster in scATAC-seq data: (A) B cell, (B) CD14+ Monocytes, (C) CD4+ naïve T, (D) CD4+ T, (E) CD4+ T 2, (F) CD8+ T, (G) CD8+ T 2, (H) NK, and (I) Plasma B cell. For each annotated cluster, the motif detection was performed via SnapATAC interface, where the motif sequences enriched around the peaks called by MACS2 are estimated using HOMER.

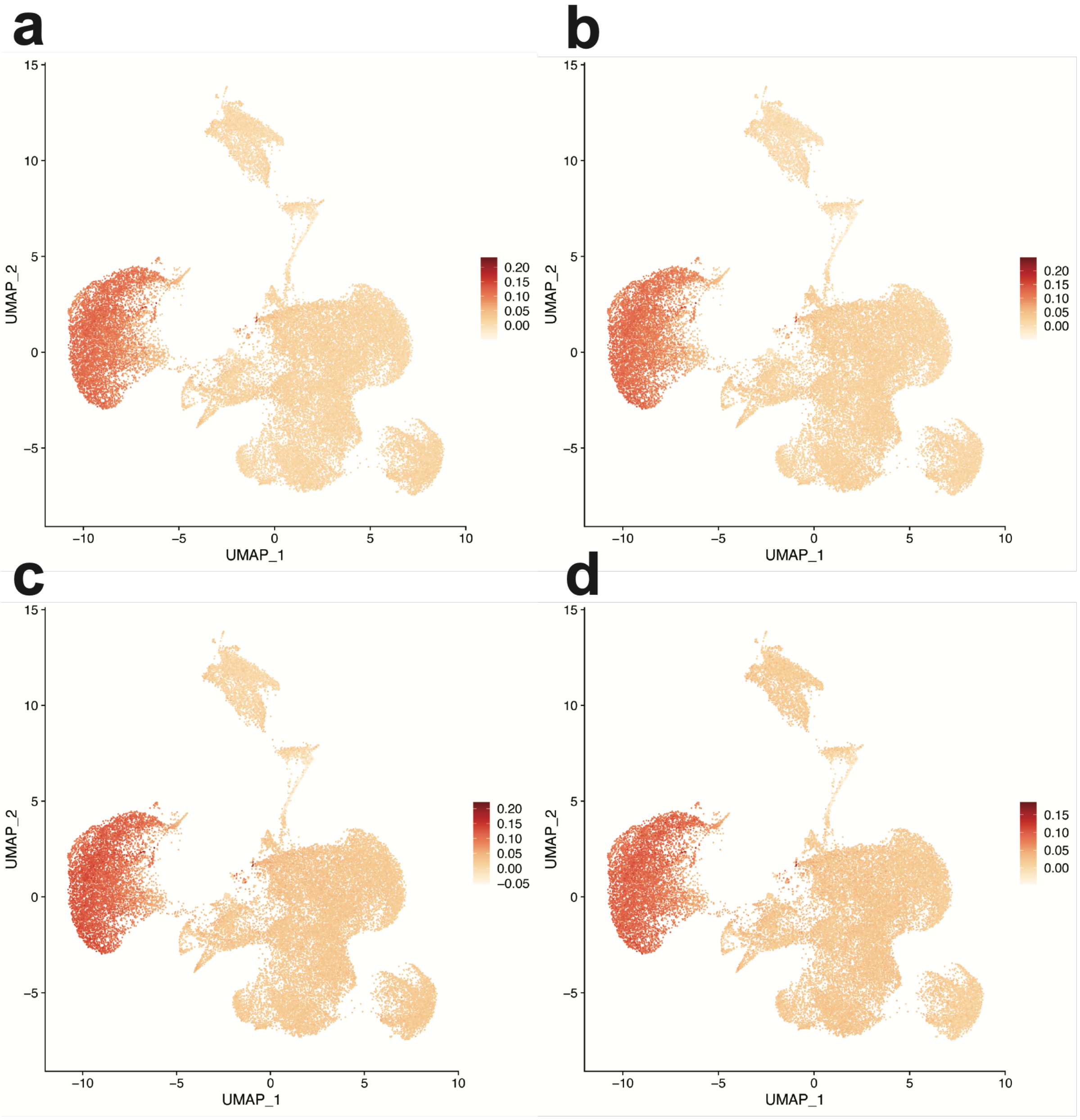


**Fig. S15.**

Module scores of the predictive genes for $\beta$. Aggregated expression level is computed for the predictive genes specific for 16-90 $\beta$ at (A) $t_1$, (B) $t_2$,(C) $t_3$, and (D) $t_4$ (single-cell pseudo bulk data) based on the average ranking of training data, which are bulk transcriptome at two other time points (A-C), or $t_1$ and $t_2$ (D).
